# Experimental evidence for female choice in an angiosperm

**DOI:** 10.64898/2026.07.28.741183

**Authors:** T. Chenin, F. Rousset, E. Barbot, A. Mignot, J. Tonnabel

## Abstract

Female choice is a central process of sexual selection that has shaped numerous phenotypes in animals. It may operate broadly across sexually reproducing organisms, including plants, not only because sex differences in sexual selection arise from anisogamy, wich refers to the unequal investment in gametes between sexes, but also because it can emerge from simple variation in female reproductive morphology. Here, we provide the first empirical test of cryptic female choice in an angiosperm by examining whether pollen and pistil traits jointly influence paternal fertilization success. Using experimental pollen competition, paternity analyses, and trait measurements, we show that pistil traits can bias paternity toward pollen donors with specific pollen traits. These results demonstrate that cryptic female choice operates in plants, paralleling mechanisms described in animals.

## Introduction

Sexual selection is widely recognized as a powerful evolutionary force shaping diverse reproductive strategies in animals [1]. Sex-specific sexual selection typically arises from anisogamy – the difference in gamete size and number between sexes – where females produce fewer and larger gametes than males [2–4]. This numerical asymmetry underpins a foundational prediction of sexual selection theory: male reproductive success is expected to be limited by access to sexual partners and their gametes, whereas female reproduction should rather be constrained by their ability to allocate resources to offspring production [2–4]. Consequently, anisogamy promotes male-male competition for access to the limiting female gametes, while females potentially exert selective pressure through female choice, biasing paternity towards particular males. Sexual selection theory therefore, predicts its universal operation across all sexually reproducing anisogamous organisms, including plants [2,5,6]. However, research on sexual selection in plants has long been hindered by conceptual barriers, stemming from their lack of cognitive ability, their predominantly hermaphroditic reproductive system, and the limited sexual dimorphism observed in the rarer dioecious species [7–9]. Despite these initial obstacles, recent studies have provided experimental evidence supporting core predictions of sexual selection theory, demonstrating that sexual selection for access to mates can act both in hermaphroditic animals [10,11] and plant species [5,12–14], thereby removing the conceptual barriers that once excluded plants from this theoretical framework.

Although sexual selection is nowadays accepted as a potential force shaping plant traits, the potential for female choice – a central pilar of sexual selection in animals – has been surprisingly overlooked. This gap likely reflects that the possibility for female choice to extend beyond pre-mating cognitive partner selection to include post-mating morphological mechanisms was only considered secondarily, even in animals [15,16]. Such cases, termed cryptic female choice, occur when variation in female reproductive morphology biases fertilization toward sperm exhibiting particular traits [15–17]. An iconic example is found in *Drosophila*, where the evolution of gigantic sperm is driven by selection from females with larger sperm-storage organs [18]. These findings demonstrate that the core assumptions of female choice models – Fisher’s runaway and good-genes processes – can be fulfilled by simple morphological female traits that directionally biases paternity toward a male morphological trait, without requiring cognition, thus extending naturally to plants [6]. Under

Fisher’s runaway model, female choice can evolve because mate choice alleles become associated with alleles for the preferred male trait in sons of choosy females, thereby gaining a transmission advantage and possibly offsetting the costs of preference [19,20]. The good-genes model additionally posits that the preferred male trait reflects overall genetic quality, enabling choosy females to produce sons that are fitter not only because they are more attractive [21]. Although developed with an animal focus, these models are not animal-specific and can theoretically apply to plant reproduction. In plants, the post-pollination phase – during which male and female gametophytes physically interact – offers candidate pistil traits for paternity biases in ways analogous to female genital tracts, with potential to parallel cases of cryptic female choice in animals [6].

The lack of investigation of cryptic female choice is surprising, given that decades of pollination and molecular research have established the conditions for its evolution during the post-pollination phase [6]. In most species, pollen deposition on stigmas often far exceeds ovule numbers, setting the stage for pollen competition [22]. Multiple paternity within fruits confirms that pollen donors compete for ovule access in natural populations [22]. Paternal fertilization success after artificial pollination is consistently linked to pollen traits, such as pollen size [23,24], germination rate [25,26] and pollen-tube growth rate [27,28]. Furthermore, many sporophytically expressed genes are active in gametophytic tissues [29], rendering post-pollination interactions plausible arenas for good-genes processes. Analogously to sperm length in animals [18], pollen-tube growth rate can predict offspring performance, thus forming an honest indicator of genetic quality [30–32]. Moreover, molecular studies have revealed that pistil tissues are far from passive: they actively influence pollen-tube growth and guidance [33], providing molecular mechanisms on which selection could act to favour the paternity of certain males. Yet, variation in paternal fertilization success in plants has been studied almost exclusively through the lens of pollen competition, with a lack of experimental tests of directional cryptic female choice.

Here, we experimentally tested for female choice during the post-pollination phase in the hermaphroditic angiosperm *Brassica rapa.* We tested the hypothesis that pistil traits determine the outcome of competition among pollen donors, effectively biasing paternity toward donors producing specific pollen traits [6,34]. Longer pistil structures (stigma, style, and ovary) may select for faster-growing pollen tubes by increasing the distance theses tubes must travel, and wider stigmas may select more competitive pollen traits by increasing the number of competing pollen grains. We manually pollinated recipient plants with pollen from five paternal donors, to create pollen competition arenas. Both parents and offspring were genotyped for paternity analysis, and pollen and pistil traits were exhaustively measured and reduced to uncorrelated components reflecting the distance to reach the ovules (style and ovary lengths) and the surface available for pollen competitor accumulation (stigma area). Our main experiment involved 70 pollen recipient plants, 70 pollen donor plants, and 2,122 genotyped seeds. It was complemented by a second experiment using fewer parents but extensive within-parent trait measurements to assess trait repeatability and to provide an independent, albeit lower-powered, test of cryptic female choice. We found evidence for female choice processes, including two qualitatively distinct patterns: unidirectional choices pattern, as commonly assumed in models of sexual selection, and an opposite-directional choices depending on the female trait.

## Results

Validating core assumptions of female choice in plants, we show that models with interactions of pollen and pistil traits jointly explain variation in paternal fertilization success under pollen competition, involving multiple uncorrelated pollen-pistil trait pairs (Table 1; Figs 1 and 2). In models without effects of pistil traits, pollen donors with higher *in vitro* germination rates tended to fertilize more ovules (Tables S1 and S2), confirming germination rate as a competitive pollen trait [25,26].

**Fig 1.**
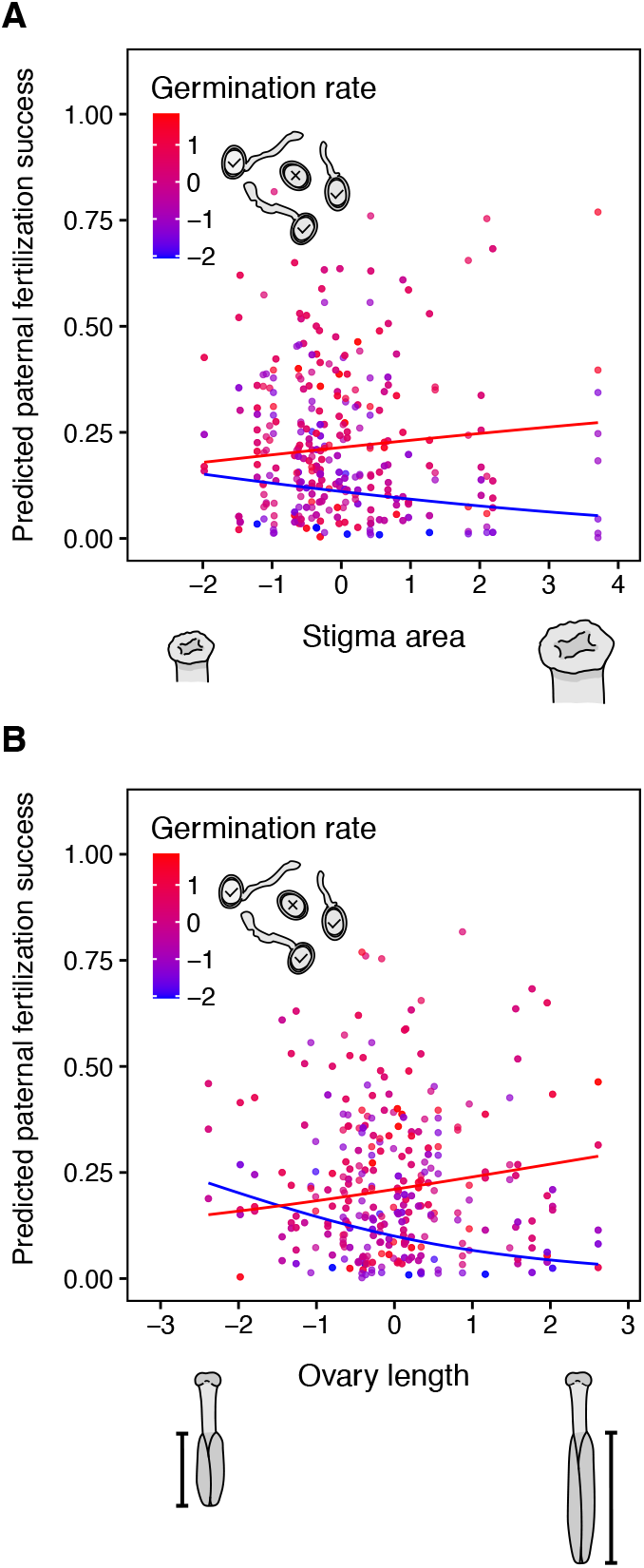
Directional effects of pollen-pistil trait interactions on predicted paternal fertilization success. Points show predicted fertilization success of pollen donor plants across pistil trait values, with germination rate values indicated by a blue-to-red gradient. Blue and red lines indicate model predictions for low (0.1 quantile) and high (0.9 quantile) pollen trait values, respectively. Panels display significant interactions between (**A**) pollen germination rate and stigma area and (**B**) pollen germination rate and ovary length. The model included germination rate, pollen size, and tube growth rate as main effects, and their interaction with stigma area, style length and ovary length. Relative pollen production was included as a covariate, and pollen donor plant identity and source population were included as random effects. All pollen and pistil traits were standardized and correspond to predicted values extracted from individual-level random effects of generalized linear mixed-effects models (GLMMs) that corrected for pollen density and other potentially confounding experimental factors included as random effects.

**Fig 2.**
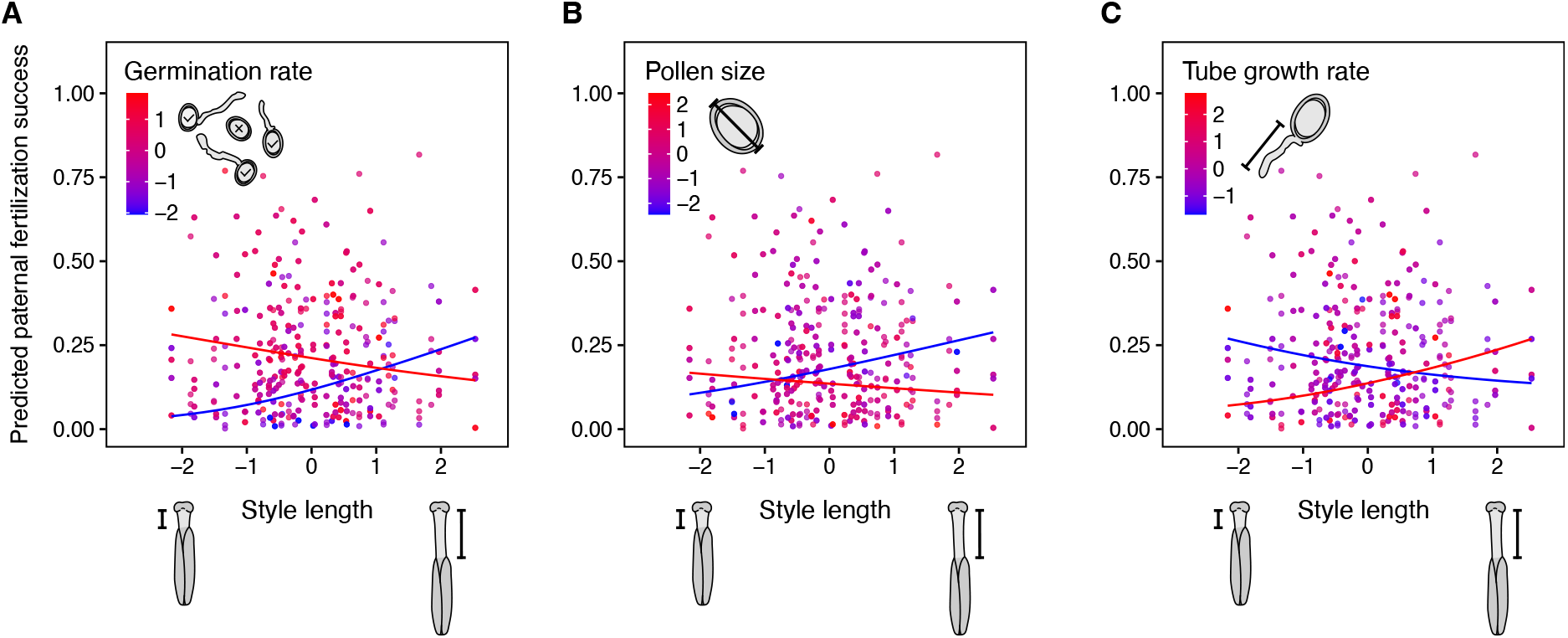
Opposite-directional effects of pollen-pistil trait interactions on predicted paternal fertilization success. Points show predicted fertilization success of pollen donor plants across pistil trait values, with pollen trait values indicated by a blue-to-red gradient. Blue and red lines indicate model predictions for low (0.1 quantile) and high (0.9 quantile) pollen trait values, respectively. Panels display significant interactions between style length and (**A**) germination rate, (**B**) pollen size, and (**C**) tube growth rate. The model included germination rate, pollen size, and tube growth rate as main effects, and their interaction with the focal pistil trait. Relative pollen production of competitors was included as a covariate, with pollen donor plant identity and source population included as random effects. All pollen and pistil traits were standardized and correspond to predicted values extracted from individual-level random effects of generalized linear mixed-effects models (GLMMs) that corrected for pollen density and other potentially confounding experimental factors included as random effects.

**Table 1.**
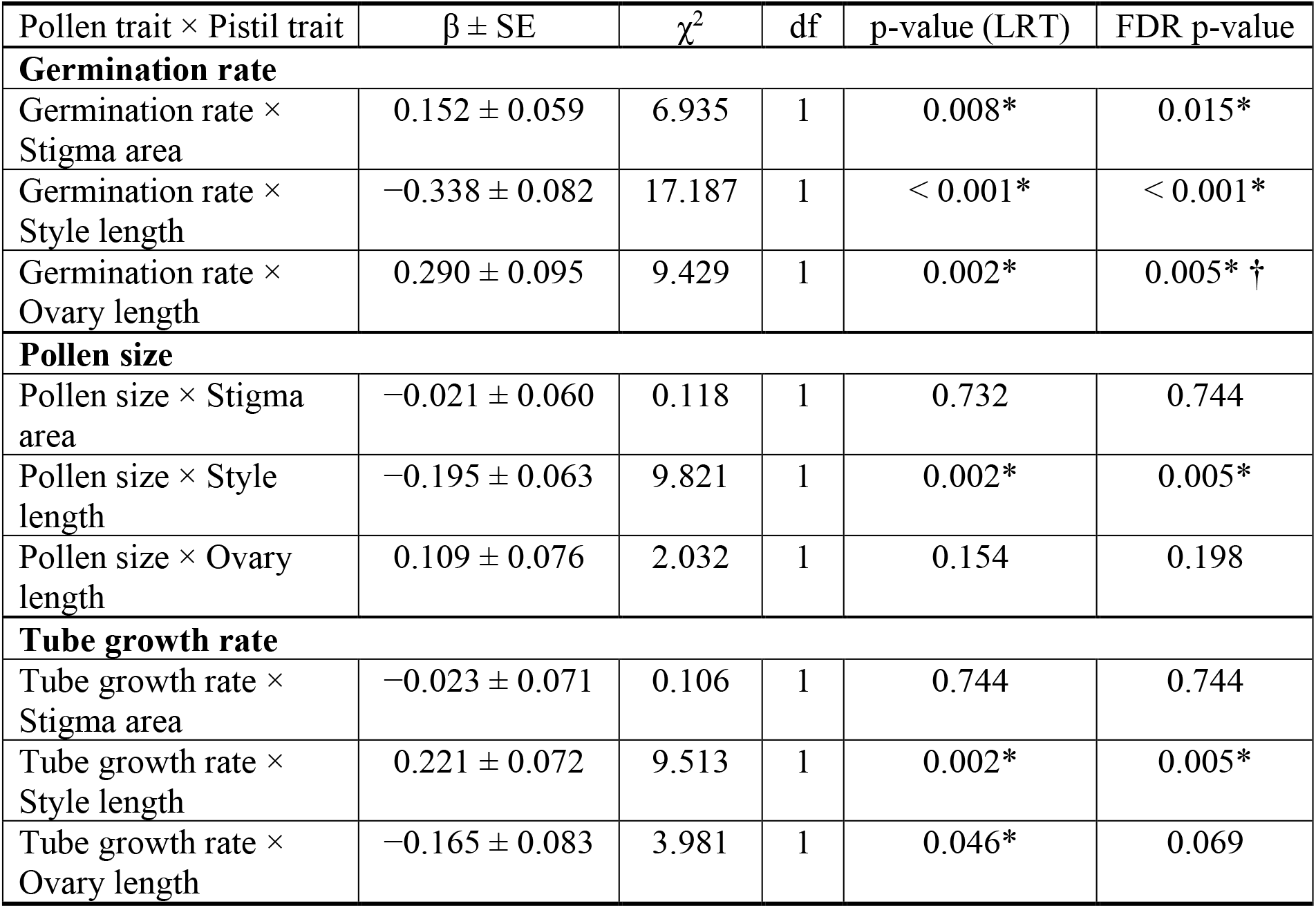
Estimated effects of pollen by pistil traits interaction on paternal fertilization success.

### Evidence for directional post-pollination female choice

For some trait combinations, female choice was directional, *i.e.* in the same direction for low and for high values of the female trait. In particular, the advantage of high germination rate was amplified in pistils with larger stigma areas (Table 1; Fig 1A) and in pistils with longer ovaries (Fig 1B), a trait statistically independent of other pistil dimensions (except for ovary width; Spearman’s ρ = 0.41, p < 0.001; Fig S1A). Thus, pollen with higher germination rate achieved greater fertilization success, particularly when competing within wider pistils or those with longer ovaries. Potential confounding effects on this relationship (and others documented below) were accounted for: pollen density influenced *in vitro* germination rates negatively or positively, respectively, in our main and complementary experiments, illustrative of the dual effects pollen density often found in other plant species [35]; Tables S3 and S4), whereas fertilization success was not explained by variation in pollen production, calculated as the number of pollen grains produced by a given pollen donor relative to the number produced by other competitors (Tables S1 and S2; see also [36]). Moreover, our statistical models also included individual random effects to account for donor non- independence among competition arenas (see Material and Methods). These cryptic female choice patterns were broadly robust in a complementary experiment with fewer parental plants (Tables S6 and S7), confirmed by Fisher’s combined probability test revealing a significant global effect of pollen-pistil interactions across both experiments (Tables S8 and S9). Together, interactions between pollen germination rate and independent components of pistil morphology (Table 1; Fig 1), as well as additional pollen-pistil trait interactions (Table 1; Tables S5–S7), indicate that cryptic female choice operates at multiple stages of pollen- pistil interaction. Overall, our results demonstrate that simple variation in pistil morphology can bias fertilization toward pollen with distinct germination rate, consistent with directional female choice.

### Evidence for opposite-directional selection on pollen traits

Our results also revealed a second form of female choice driven by pistil morphology that can be described as opposite-directional, whereby opposite ends of a pollen trait distribution are favored by opposite extremes of a pistil trait. Such pollen-pistil interactions were detected for multiple trait combinations, involved all pollen traits measured, and were qualitatively distinct from the directional patterns reported above. For instance, pollen with higher germination rates fertilized more ovules in pistils with shorter styles, whereas pollen with lower germination rates was favored in pistils with longer styles (Fig 2A). An identical pattern was observed for pollen size (Fig 2B), despite the absence of correlation between pollen size and germination rate (Fig S2A; Spearman’s ρ = −0.20, p = 0.307), indicating independent axes of female choice. Style length also modulated selection on pollen tube growth rate: faster- growing tubes were favored in longer styles, whereas slower-growing tubes achieved higher success in shorter styles (Fig 2C). Although pollen tube growth rate and pollen germination rate were positively correlated (Fig S2A), the contrasting effects of style length on these traits demonstrate distinct female choice processes. The opposite-directional results of the main experiment were broadly robust in the experiment involving fewer parental plants, though some were significant only under simpler models (models with a single pollen-pistil interaction by contrast to models with multiple interactions; Tables S6 and S7; see Fisher’s test results; Tables S8 and S9). An additional opposite-directional interaction, detected only in the complementary experiment, involved larger stigmatic area which favored smaller pollen grains and faster-growing tubes, while smaller stigmas favored larger pollen grains and slower-growing tubes (Fig S3; Tables S6 and S7). Consistent with these findings, pollen tube growth rate and pollen size did not influence fertilization success overall in the model without pistil trait effects (Tables S1 and S2).

Are measured pollen and pistil traits individual plant-level characteristics, as required for their potential evolution through sexual selection? All traits involved in the detected patterns of cryptic female choice were significantly repeatable among flowers within individuals (Fig 3; Tables S10 and S11) and, for pollen traits measurable across multiple stamens, within flowers (Fig 3A; Table S10). In the experiment with repeated measurements on a limited number of parental plants, estimated repeatabilities ranged from 0.259 to 0.462 for pistil traits (Fig 3B; Table S11), from 0.222 to 0.319 for pollen dynamic traits, and 0.259 to 0.678 for pollen size (Fig 3A; Table S10), all quantified at the plant scale. These values – estimated after controlling for all potential experimental confounding effects – indicate that traits implicated in cryptic female choice reflect consistent plant-level characteristics, a necessary condition for their evolution as repeatability sets the upper limit of heritability.

**Fig 3.**
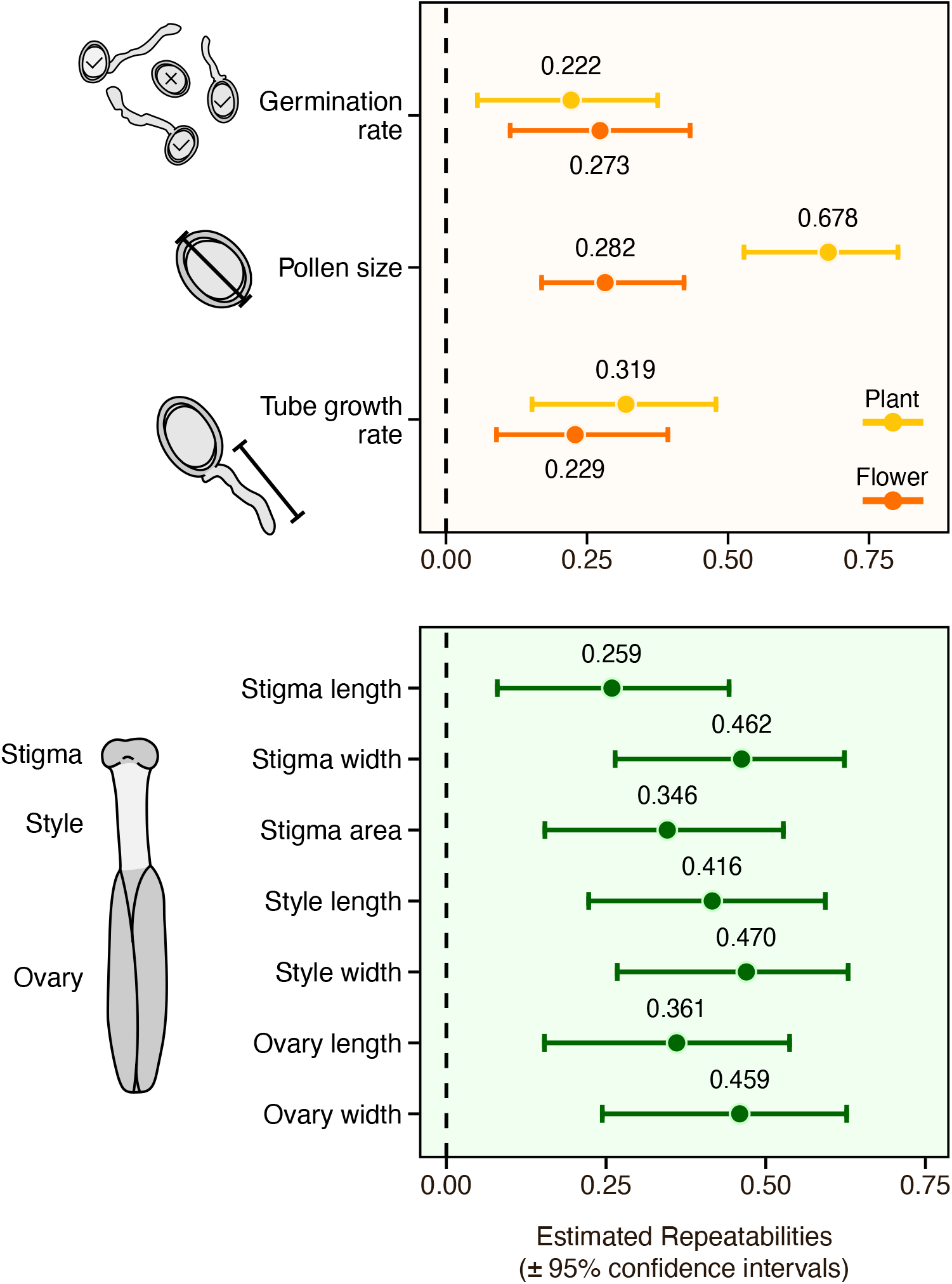
Estimated repeatabilities (± 95% confidence intervals) for pollen (A) and pistil traits (B). Repeatability represents the proportion of total trait variation attributable to differences among plants relative to within-plant variation. Dots indicate mean repeatability estimates across 1,000 simulations, and bars represent the corresponding 95% confidence intervals. Standardized trait proxies extracted from GLMMs correcting for relevant experimental covariates were used as response variables, with plant identity included as a random effect and flower identity additionally included for pollen traits. Gaussian error distributions were assumed, and pollen tube growth rate was square-root transformed. Repeatability estimates are represented on the original scale.

Importantly, the observed patterns of cryptic female choice are unlikely to result from genetic self-incompatibilities characteristic of *Brassica* species. Most pollen-pistil interaction effects on fertilization success remained robust in sensitivity analyses excluding pollen donor plants that sired less than five percent of seeds under competition, at least in simpler models considering one pollen trait at a time (Table 1; Tables S12–S15). Moreover, the proportion of pollen donors failing to fertilize any seeds in competition experiments (19.52%) greatly exceeded the near-zero rate of incompatible crosses measured in experiments using the same *B. rapa* line. This indicates that low fertilization success reflects poor competitive performance rather than incompatibly at the self-incompatibility locus, confirming that the detected cryptic female choice involves interactions among compatible mating partners, as assumed by classical female choice models.

## Discussion

We provide experimental evidence for female choice during the post-pollination phase in an angiosperm, demonstrating that female morphological traits directly bias paternity toward specific pollen traits – a core requirement of classical female choice models. These findings parallel the widespread occurrence of female choice in animals, which involves not only behavioural traits [37–39] but also the morphology of female reproductive tracts selecting sperm traits [15–18]. In plants, the closest evidence to date comes from a study in wild radish, where seeds deeper in ovaries are preferentially aborted under water deprivation, potentially favoring pollen tubes that grow faster [40]. However, this case lies at the interface between post-pollination sexual selection and post-fertilization natural selection and remains indirect, as it is not linked to explicit pollen and pistil traits [40]. By contrast, our study combines competitive pollination and exhaustive measurements of pollen and pistil traits to provide direct evidence that pistil morphology mediates selection on pollen traits. Importantly, we show that multiple independent pollen-pistil trait pairs shape fertilization success at different stages of the pollen tube journey. We identify two qualitatively distinct forms of cryptic mate choice in *B. rapa*: one aligning with classical female choice models, involving consistent directional effects of pistil traits on male performance, and another in which opposite ends of a pollen trait distribution are favored by contrasting pistil phenotypes. Together, these findings reveal that pollen-pistil interactions are more diverse and mechanistically complex than previously recognized [6,34], and that female reproductive morphology actively shapes competitive fertilization outcomes.

Several pollen-pistil interaction patterns underlying cryptic female choice support our functional hypothesis that female traits exacerbating pollen competition select for pollen competitive traits. Pollen donors producing pollen with higher germination rates – the primary determinant of fertilization success and a confirmed competitive trait in *B. rapa*, as in other species [25,26] – were consistently favored in pistils with larger stigmas and longer ovaries. These two pistil traits were expected to intensify pollen competition by increasing respectively the number of competing pollen grains and the distance pollen tubes must travel to reach the ovules. Some opposite-directional female choice patterns also aligned with this idea, as faster-growing pollen tubes were favored by both longer styles and larger stigmatic areas, and conversely slower-growing tubes were favored by opposite female traits. Additional comparable patterns were reported only in our less replicated experiment, with larger stigmatic areas favoring faster-growing pollen and smaller pollen grains for most of the pistil distribution, potentially reflecting a numerical competitive advantage under exacerbated pollen competition. Accordingly to female choice models, our reported paternity biases at the micro-evolutionary scale should foster genetic correlations between pollen and pistil traits, and drive patterns of male-female trait coevolution across taxa, as observed in animals [15–18]. Such genetic correlations have been reported in our study species between pollen size and style length [24] converging with recent patterns of male-female trait coevolution found in a meta-analysis spanning 89 plant families [41], but this specific trait association was not supported by our data.

Our results reveal more complex pollen-pistil interactions than predicted by our simple hypothesis linking competitive traits to intensified competition. In particular, longer styles favored pollen with lower germination rates. Moreover, several opposite-directional patterns observed across parts of the pistil are difficult to interpret from an adaptive perspective, as it is unclear why female reproduction would benefit from favoring slower-growing pollen tubes or pollen with lower germination rate. Such results may reflect different biological processes. First, purely Fisherian processes may arise from any pair of male-female traits, without requiring selection to initially target the most competitive male traits. Second, reduced pollen tube growth may be correlated with other fitness components, such that the observed paternity bias pattern may ultimately reflect a good-genes process. A limitation of our study is that we did not assess the relationship between pollen traits and fitness, a core assumption of good-genes models [21], already validated for sperm traits [18,42,43]. However, evidence suggests that pollen competitive traits values – rather than opposite values favored in some of our reported female choice patterns [32,44,45] – and intensified competition among pollen grains [36,46] can enhance progeny vigor. Third, reduced pollen tube growth may trade-off with other aspects of pollen performance not measured here. In particular, it may reflect response of pollen to the active role of pistil tissues in both pollen tube growth and guidance [33,47]. Complex phenotypic changes in pollen tubes are commonly induced by interactions with pistil tissues [48,49] – processes not been fully captured by *in vitro* pollen tube assays or morphological traits. Consistent with this idea, most of the highlighted patterns of paternity biases that deviate from our functional hypothesis involve style length, the pistil region known to assist heterotrophic pollen tube growth [33].

While our documented cases of directional cryptic female choice align with classical models of female choice, the opposite-directional patterns observed here may reflect alternative female morphological strategies with dual mating preferences coexisting in the study population, as reported in some animal taxa [17,50]. These patterns also remain compatible with good-genes processes under certain conditions. Importantly, opposite-directional patterns of paternity bias could still drive directional evolution of pistil and pollen traits if one side of the pattern is associated with higher fitness. In this context, pistil traits favoring lower-fitness pollen may represent costs of choice, with variation driven by individual differences in genetic quality [51]. Our study population originates from an experimental line established from numerous natural populations and retains high genetic and phenotypic variability, including in the traits studied here [24,52]. Consequently, natural populations with narrower variation may express only a subset of these opposite-directional patterns, more closely matching Fisher’s runaway process. Finally, these opposite-directional female choice patterns are not the mere result of genetic self-incompatibility in *B. rapa*: the rate of plants failing to fertilize any seed far exceeds that expected from incompatible monogamous crosses, suggesting the involvement of compatible but poorly competitive pollen donors. Our results remain robust after excluding these pollen donors, a conservative scenario given their likely compatibility with the pollen recipient. Our results therefore reflect mate choice mechanisms distinct from widespread genetic self-incompatibility systems – a form of mate choice that differs from female choice models [6].

We provide experimental evidence that pollen and pistil traits interact to shape fertilization success, demonstrating multiple cases of female choice in a plant species, consistent with classical sexual selection theory. Our results underscore the importance of considering female choice as a selective force, paralleling the numerous empirical findings in animals. Building on this validation of basic female choice mechanisms, future research should test predictions of Fisherian and good-genes models and quantify the evolvability of female choice, beyond the repeatabilities reported here, which represent the upper limit of trait evolvability. Whether these cases of female choice shape pollen and pistil trait evolution across angiosperms, and whether these morphological traits act independently of pollen-pistil molecular interactions, remain open questions that could be addressed by integrating plant molecular studies with classical empirical tests of the theory as performed in animals, including quantitative genetics and experimental evolution [6,53–55].

## Material and Methods

### Biological material

*Brassica rapa* (Brassicaceae) is an herbaceous, hermaphroditic, insect-pollinated plant species widely distributed throughout Europe and North America [56,57]. Our experiment was carried out with seeds derived from the Wisconsin Fast Plants™ lineage [52] (WFP™; Carolina Biological Supply Company, Burlington, North Carolina, USA). This line was established from seeds harvested from multiple wild cabbage populations across North America. Following this collection, the *B. rapa* Fast Plant line underwent selection for short life cycle (∼8 weeks) and high fecundity while maintaining substantial phenotypic and genetic variability [52]. *B. rapa* possesses a self-incompatibility genetic system known to prevent self-fertilization in the *Brassica* genus [58,59]. However, moderate levels of selfing have been reported in this species, including in the study line used here [56]. Flowers produce four long (superior) and two short (inferior) stamens that are spatially separated from the pistil, a form of herkogamy that further limits self-pollination. Together, these characteristics facilitates the manipulation of the number and identity of pollen donors during hand-pollinations, and justified treating pollen competition to be mainly allogamous. Seeds of *B. rapa* Fast Plant purchased from the University of Wisconscin [52] were used to establish five independent lines maintained under strong polygamy for three to four generations. At each generation, 85 parental plants were grown, and hand-pollinated using a pollen mix collected across all plants in the experimental population and with from twelve stamens harvested per plant. The resulting seeds formed five “source populations” used to generate pollen recipients (maternal plants) and pollen donors (paternal plants) in our experiment.

Some pollen and pistil traits examined here as candidates for cryptic female choice have been shown to exhibit substantial genetic variance in the *B. rapa* WFP™ [52]. Specifically, artificial selection experiments have demonstrated that pollen size is highly heritable, with additive genetic effects accounting for more than 30% of its phenotypic variation [24]. Furthermore, selection for larger pollen size was accompanied by increased style length [24], indicating a positive genetic correlation between these traits. This correlation is consistent with predictions of Fisherian runaway and good-genes models of sexual selection [19–21].

### Overview of the conducted experiments

Our main experiment tested cryptic female choice in *B. rapa* by creating multiple pollen competition arenas, each consisting of five pollen recipients manually pollinated with a mixed pollen sample collected from five pollen donors. Cryptic female choice was evaluated by extensively measuring pollen and pistil traits, genotyping seeds to assess paternal fertilization success, and testing whether fertilization outcomes were jointly determined by pollen and pistil traits. A complementary experiment replicated the main design with fewer competition arenas but substantially increased trait measurement per individual. This experiment thus provides a lower-powered temporal replication of the main experiment while informing the evolutionary potential of putative cryptic female choice traits, as trait repeatability sets an upper bound on heritability [60].

### Experimental design

Our main experiment consisted of mating groups composed of five pollen donors whose pollen was artificially competed on pistils of five pollen recipient plants. It was conducted in January 2023 in the greenhouses of the CNRS experimental platform in Montpellier, France. Seeds from the source populations were sown in two temporal blocks spaced one week apart to ensure a continuous supply of sexually mature plants. Plants were grown individually in 0.7 L pots filled with a sterilized soil mixture and maintained under standardized greenhouse conditions (continuous light, 25°C). Plants were placed in mesh cages and physically spaced apart to prevent physical contact and insect-mediated pollination. For each “mating group”, five pollen donors and five pollen recipients were randomly selected within a temporal block, subject to standardized floral age criteria. Pollen donor plants bore at least one freshly opened flower with non-dehiscent stamens, while recipient plants had at least two flowers about to open, to minimize pollen loss due to anther dehiscence and undesired pollen deposition prior to hand-pollination, respectively. Focal flower, hereafter termed “donor” and “recipient”, were marked and identified individually the evening before pollination. Both recipient and donor plants in each mating group originated from different “source populations”. In total, fourteen mating groups were established in our main experiment, comprising 70 pollen recipients and donor plants (see below for a description of the complementary experiment).

Hand-pollinations of multiple mating groups were conducted on the same day and are hereafter referred to as “pollination sessions”. On the morning of each session, two non-dehiscent stamens were collected from donor flowers on each of the five pollen donors. Stamens were placed in Petri dishes under a lamp to accelerate dehiscence for three to four hours. Once dehisced, pollen was mixed under a binocular microscope using a brush and then used to pollinate recipient flowers on the pollen recipients. The pollen mixture was applied to the stigmas, thoroughly saturating them. This pollination protocol was designed to minimize selective processes during the pre-pollination phase that could influence paternal fertilization success. Because the number of pollen grains deposited greatly exceeded the number of ovules, pollen donors effectively competed for fertilizing ovules.

Pollen deposition on pistils was standardized by manually pollinating one recipient flower per pollen recipient in a randomized order, followed by a second flower in the reverse order. The brush was reloaded between the two passages to homogenize pollen deposition quantities across the pollination sequence and changed to minimize pollen contamination among pollen recipient caused by inadvertent pollen transfer on the brush. Despite this precaution, a fraction of paternities was attributed to pollen recipient plants, likely due to pollen transport between successive pollinations (see below). These fertilization events between pollen recipients probably did not prevent competition to occur among the five pollen donors studied, but reduced the number of offspring available for comparing competitive outcomes. Accordingly, genotyped seeds assigned to pollen recipient plants within mating groups were excluded from subsequent analyses. The pollen mix from the different pollen donors was applied in excess to ensure overall stigmatic saturation; however, the relative contribution of each pollen donor was not standardized beyond the fact that each pollen donor contributed two stamens to the pollen mix. Therefore, the pollen mixture applied may reflect differences in pollen production in the two stamens collected per donor plant. Because pollen quantity deposited on stigmas can be a key determinant of paternal fertilization success in mixed hand-pollinations [36,61,62], and because we did not directly control donor-specific pollen deposition, we quantified pollen production among pollen donors and incorporated this variation into our statistical analyses.

### Pollen traits measurements on pollen donor plants

We collected two distinct superior stamens from each donor flower (*n* = 70) to estimate pollen traits and pollen production per pollen donor. Pollen traits were measured from dehiscent stamens of freshly opened donor flowers, whereas pollen production was estimated from non-dehiscent stamens collected immediately after flower opening to prevent pollen loss (see below). Pollen germination assays were conducted immediately after stamen collection and separately for each stamen collected from each pollen donor (*n* = 70). Fresh pollen was extracted using an insect needle and deposited onto a microscope slide containing 20 µL of pollen growth medium (PGM) optimized for *B. rapa* and adapted from previously described protocols [63,64]. To prepare 100 mL of PGM, we diluted 10 g of sucrose and 3 g of PEG 6000 in 98 mL of distilled water, then added 0.5 mL of a 0.05 g/10 mL (0.5%) H₃BO₃ solution, 0.5 mL of a 1 M/10 mL CaCl₂ solution, and 2.55 µL of a 1 M/10 mL KH₂PO₄ solution. Immediately after pollen deposition, slides were placed in an incubator at 20°C inside Petri dishes containing a 3.5 x 7 cm cotton pad soaked with 2 mL of water to prevent dehydration and maintain optimal humidity for pollen germination and pollen tube growth.

Images of pollen tube growth were acquired using an optical microscope with a 10× objective (Olympus CX43; Olympus Corporation, Tokyo, Japan). Images were acquired after one, three, and five hours of growth on the medium. For each slide and time point, five images were captured from different areas on the slides to sample a large number of pollen grains while avoiding duplicate measurements. For pollen donors belonging to three mating groups, images were acquired using a 5× microscope objective instead of a 10× objective; this difference was accounted for by including objective magnification as a random effect in subsequent statistical analyses. Pollen trait measurements were obtained by processing images using the ImageJ software [65]. On each image, we measured traits reflecting pollen growth dynamics: (i) germination rate, estimated by classifying and counting pollen grains as germinated or non-germinated, (ii) pollen size, measured as the diameter of three germinated and three non-germinated pollen grains, and (iii) pollen tube growth rate, quantified as pollen tube length after a standardized growth period in the medium, measured on the three germinated pollen grains selected for pollen diameter measurement and two additional germinated grains. Pollen grains from both categories were selected using a semi-automated protocol based on randomly generated coordinates. For each image, the pollen grain nearest to a given coordinate was selected for measurements. The first three coordinates targeted non-germinated pollen grains, while the remaining five random coordinates were used to identify germinated pollen grains for growth dynamics measurements.

Following common practice, a pollen grain was considered germinated when pollen tube length was at least equal to the pollen diameter [66,67]. A small proportion of pollen grains were classified as sterile because they were markedly smaller and failed to produce pollen tubes. In addition, some pollen grains could not be reliably classified due to poor image quality caused by suboptimal microscope focus or pollen aggregation, and were therefore labeled as indeterminate. For each slide and time point, the five images were randomly assigned to three measurers, whose identity was included as a random effect in all subsequent analyses. Across the three standardized time points and five images per time point, we measured on average (± SD) 28.31 ± 10.21 non-germinated pollen grains, 28.67 ± 10.78 germinated pollen grains, and 46.24 ± 17.05 pollen tubes per pollen donor. One individual was excluded from the statistical analyses because all pollen grains were sterile, preventing trait measurements. Because germinated and non-germinated pollen grain diameters were highly and positively correlated in both experiments (main experiment: Spearman’s ρ = 0.57, p < 0.001; complementary experiment: Spearman’s ρ = 0.80, p < 0.001), we report only results for germinated pollen grain diameter, referred to as “pollen size” in the main text and in the figures. Germination rate was estimated by counting germinated (367.10 ± 215.76), non-germinated (487.35 ± 338.49), and sterile pollen grains (48.54 ± 100.80) per pollen donor on average (± SD). Although our protocol minimized variation in pollen deposition across collected stamens, we did not precisely control the number of pollen grains deposited on microscope slides. Because pollen density affects pollen growth dynamics [35,68,69], the total pollen grain count per image, regardless of pollen category, was included as a covariate in all subsequent statistical analyses (see below).

### Pollen production on pollen donor plants

Pollen preparation for counting was inspired by a previously described protocol [70] and adapted for *B. rapa*. Each collected stamen per pollen donor was stored individually in a 1.5 mL tube at –80 °C prior to pollen extraction. Samples were incubated in 200 µL of sulfuric acid for 48 hours to digest residual stamen tissue. Tube contents were then crushed with a glass rod to release pollen from anther tissues. Samples were subsequently treated with 100 µL of distilled water and subjected to ultrasonic treatment to dispersed aggregated pollen grains. Tubes were centrifuged at 820 rpm for two minutes, and the supernatant was discarded, leaving a pollen pellet. A second ultrasonic treatment was performed after adding 1,000 µL of pure ethanol, followed by centrifugation at 14,000 rpm. The final pollen pellet was resuspended in 400 µL of distilled water and stored at 3–6 °C to prevent evaporation.

Pollen production per pollen donor was estimated by counting pollen grains on Mallassez slides (5 × 5 grid) under an optical microscope (Olympus CX43; Olympus Corporation, Tokyo, Japan) with a 10× objective. For each collected stamen, two successive 10 µL aliquots of the pollen suspension were deposited onto the slide and pollen was counted in these two technical replicates. We applied a standard cell-counting protocol, whereby pollen grains overlapping the left or bottom borders of a grid cell were excluded to avoid double-counting in a given solution volume. In rare cases, pollen grains aggregated on the medium, forming clumps containing variable numbers of pollen grains (ranging from approximately ten to a maximum of one hundred). When this occurred, the number of pollen grains was visually estimated based on the aggregate size. In total, we counted nearly 5,800 pollen grains across 65 collected stamens.

For all collected stamens, the pollen grain pellet unintentionally dried before the final resuspension step, which likely increased pollen aggregation and negatively affected counting accuracy. We therefore optimized the protocol to improve pollen grains separation as follows: (1) two droplets of distilled water containing commercial liquid soap were added using a Pasteur pipette; (2) samples underwent ultrasonic treatment for two 10 hours periods, with sonication in the presence of three glass beads (30 Hz, 5 min) between treatments; (3) samples were centrifuged at 14,000 rpm and the supernatant was removed; and (4) the pollen pellet was resuspended in 400 µL of distilled water. Using this optimized protocol, 65 of the 70 samples were successfully recovered. To validate the modified pollen preparation protocol, we assessed its consistency with the original one. In the complementary experiment, two superior stamens were collected from two flowers on each of the 30 pollen donor plants, and pollen pellets were intentionally dried before the final suspension step (*n* = 52 stamens). Pollen production per stamen was then compared with estimates obtained using the original protocol, which involved six superior stamens collected from the same two flowers (*n* = 237 stamens). Pollen counts obtained from the two protocols and for the same pollen donor were strongly and positively correlated (Spearman’s ρ = 0.69, df = 27, p < 0.001), indicating that pollen production estimates from the main experiment reliably reflect differences among pollen donor plants.

Extensive stamen sampling across multiple flowers of the 30 pollen donor plants from the complementary experiment (228 stamens from 116 flowers) revealed that pollen production was repeatable at the plant level (R = 0.27, p < 0.001) but less so at the flower level (R = 0.09, p = 0.133; see below for calculation of trait repeatability). Thus, stamens collected from donor flowers may not accurately reflect pollen production of those used for hand-pollination. Nevertheless, pollen production did not significantly affect paternal fertilization success (Table S2), suggesting that variation in pollen production is unlikely to be a major determinant of reproductive outcomes under our mixed hand-pollination conditions.

### Pistil traits measurements on pollen recipient plants

Pistil trait measurements were conducted on two pistils collected from freshly opened, virgin flowers on the same day as the pollination session. Because pistil morphology changes over time, particularly following pollen deposition on the stigma, we standardized collection for these two pistils to freshly opened flowers to control for age and minimize the risk of sampling pistils that unintentionally received pollen grains. Flowers expected to open were marked the day before anthesis and harvested upon opening. In addition, five other pistils were sampled per pollen recipient plant from newly opened yet slightly older flowers (up to two days post-opening). These pistils were collected for a separate study that involved imaging four of the seven measured pistil traits (see below), and were therefore included in our pistil trait dataset. These additional pistils were measured from images acquired using an optical microscope (Olympus CX43; Olympus Corporation, Tokyo, Japan), whereas the other pistils were measured using a binocular microscope. This methodological difference was accounted for by including microscope type as a random effect in all subsequent analyses. Because pistil sampling dates varied among pollen recipients, sampling date was included as a random factor in the statistical analyses.

Each pistil was individually stored in a fixation solution composed of ethanol, methanol, and acetic acid (EMA, in a 1:1:3 proportion), which is known to preserve plant tissues [71]. Entire pistils were photographed in longitudinal view under a binocular microscope. Stigmas were then excised using a razor blade and photographed in apical view, with the stigma oriented upwards. Throughout the imaging process, pistils and stigmas were continuously humidified with the EMA fixation solution to prevent deformation due to desiccation. Morphological measurements were obtained from the images using ImageJ software [65] (version 2.14.0). Specifically, the lengths of the different pistil components were quantified from longitudinal images: stigma length was measured from the apex to its junction with the style; style length as the distance between the stigma and the ovary; and ovary length from the style junction to the base of the ovary (Figure 3B). Widths were measured at the midpoint of each structure. Total stigma surface area was measured from apical-view images. Stigma surface area, ovary length and ovary width were measured only in the two main pistils harvested per recipient plant.

### Collecting fruits, seed classification and germination

Four to five weeks after hand-pollination, once plants had completed their life cycle, we collected dry fruits containing mature seeds from pollen recipient plants, yielding 124 collected fruits from 140 pollinated flowers. Fruits were manually opened, and seeds were classified into four visually distinguishable and consistent categories: viable, germinated, aborted, and unfertilized. Viable seeds were spherical, with a smooth seed coat and coloration ranging from dark brown to black, and constituted the majority of collected seeds (86.8% of 2,510 seeds). Aborted seeds represented a minority and were distinguished from viable seeds by their smaller size and surface depressions, indicative of incomplete embryo development (1.0%). Unfertilized seeds differed from aborted seeds by exhibiting a transparent, soft seed coat and lacking any visible embryo (11.2%). Germinated seeds, which had developed into small plantlets within the fruit, were easily distinguishable from the other categories (1.0%).

All 2,205 collected viable and aborted seeds were sown in Petri dishes containing agar (10 g L⁻¹) to obtain sufficient DNA from the resulting plantlets for genotyping. Seeds classified as germinated were excluded from germination assays because the plantlets had died before fruit collection, preventing DNA extraction. Seeds were spaced at least one centimeter apart to minimize density effects on germination and were incubated for seven days under controlled conditions (20–23°C, 16:8 photoperiod), resulting in a germination rate of 97.3% among viable seeds. All germinated plantlets were subsequently genotyped to estimate the fertilization success of each pollen donor on each pollen recipient plant, thereby assessing pollen competitive outcomes.

### Genotyping

Leaves from both pollen donor and recipient plants were collected five to seven days after hand-pollination. A detailed genotyping protocol is provided in [72]. In brief, parental tissues were dried for 24 hours in an incubator at 35°C, and 10–12 mg of material was used for DNA extraction. We genotyped all the fresh offspring plantlets obtained from seed germination, yielding a total of 2,122 offspring. DNA extraction was performed using the NucleoMag Plant kit (Macherey-Nagel Inc., Allentown, USA). Biological samples were ground using a 3 mm tungsten bead in a tissue homogenizer (Qiagen, Hilden, Germany), incubated in a thermolyser with RNase and lysis buffer, and centrifuged. Clear lysates were transferred to deep-well plates (Thermo Fisher Scientific, Waltham, USA), followed by automated magnetic bead treatment involving three successive washes with ethanol (35%, 35%, and 80%) and using a KingFisher® workstation (Thermo Fisher Scientific, Waltham, USA). A multiplex PCR was performed using eight microsatellite markers optimized for *B. rapa* and shown to be polymorphic in the study line [72], with Applied Biosystems™ Master Mix (Thermo Fisher Scientific, Waltham, USA). PCR amplifications were carried out on an Eppendorf thermocycler (Mastercycler Nexus GSX1) under the following conditions: (i) 15 minutes at 95°C, (ii) 30 cycles of 30 seconds at 95°C, (iii) 1 minute 30 seconds at 58°C, (iv) 1 minute at 72°C, and (v) 30 minutes at 60°C. PCR products were diluted 1:120 in distilled water and both formamide and a size standard marker (GeneScan™ 500 LIZ™, Thermo Fisher Scientific, Waltham, USA) were added. Amplified fragments were analyzed by capillary electrophoresis on an ABI3500XL 24-capillary sequencer (Applied Biosystems® 3130 Genetic Analyzer). Allelic profiles were scored using the GeneMapper® software (Thermo Fisher Scientific, Waltham, USA). Genotyping was repeated for 514 offspring and 26 parental samples due to illegible allelic profiles resulting from failed amplification.

### Paternity analysis

Paternity analyses were conducted using CERVUS software [73,74]. This program estimates the likelihood that each pollen donor is the true father of a given offspring, based on the offspring’s genotype and the genotype of the known pollen recipient. The software first generates simulated parental genotypes using allele frequencies from the study population and simulates a large number of reproductive events among these parents. The resulting simulated dataset is used to calculate likelihood score distributions and to determine critical values for paternity assignment at specified confidence levels. For each offspring, a paternity confidence score (LOD score) is calculated as the log-likelihood ratio of a candidate father being the true father relative to alternative candidates. We used an 80% confidence threshold (a relaxed criterion) to assign the most likely father. Offspring with missing data at least three or more microsatellite markers were excluded from the analysis, resulting in the exclusion of 203 individuals.

Separate paternity analyses were performed for each mating group. Paternity assignment criteria were based on simulations of 10,000 hypothetical offspring, allowing for a 1% genotyping error rate estimated from observed mismatches between pollen recipient plants and offspring across both experiments. The mean mismatch rate was 0.524% across the eight markers, and the mean non-exclusion probability was 0.101 (ranging from 0.063–0.168; *i.e.* the probability that a candidate father was not excluded although he is not the true father). Pollen recipient plants were included as candidate fathers to quantify unintended paternity resulting from pollen transfer among pollen recipient plants. On average, 42.9% of offspring were assigned to pollen recipient plants, with values ranging from 13.2% to 71.9% across the fourteen independent paternity analyses. Paternity data attributed to pollen recipient plants were excluded from subsequent statistical analyses, corresponding to 723 individuals. Of the 2,122 offspring genotyped, 985 assignments were retained for the analyses in the main experiment.

### Complementary experiment protocol

A complementary experiment aimed at quantifying the repeatability – the consistency of trait measurements within individuals – of pollen and pistil traits was conducted in December 2023. This experiment also served as a replication of the main experiment with minor modifications; therefore, only deviations from the main protocol are described here. Repeatability of pollen and pistil traits was assessed using 30 pollen donors and 30 pollen recipients, which were also subjected to manual pollination to provide an independent, lower-powered test of cryptic female choice encompassing six mating groups. Each mating group underwent a second hand-pollination event using one additional donor flower per pollen donor and two new recipient flowers per pollen recipient. Hand-pollinations followed the protocol described in the main text, except that we used toothpicks instead of brushes and that we changed them after each hand-pollination to minimize undesired pollen transport between successive pollinations.

Pollen traits were measured for each pollen donor using one freshly dehiscent superior stamen from donor flowers and six additional stamens collected from two other flowers (*n* = 8), yielding a total of 232 stamens from 118 flowers across all pollen donors. Pollen production was estimated using one non-dehiscent stamens collected from the two donor flowers and two additional flower (*n* = 8), resulting in 228 stamens sampled from 116 flowers and nearly 9,500 pollen grains counted. Consequently, pollen traits and pollen production were measured on the same flower only for the two donor flowers. Pistil trait repeatability was estimated by collecting five additional virgin pistils per pollen recipient, in addition to the four pollinated recipient flowers, and all pistils were measured following our main protocol using binocular microscope. Repeated measurements from multiple stamens and pistils collected across different flowers allowed estimation of the repeatability of both pollen and pistil traits at the plant scale, and of pollen traits at the flower level. On average, estimates of pollen trait were based on 254.00 ± 40.63 non-germinated pollen grains per pollen donor, 211.83 ± 50.60 germinated pollen grains, and 329.40 ± 86.18 pollen tubes per pollen donor. Germination rate was estimated by counting on average (± SD) 3238.03 ± 1216.55 germinated, 1107.23 ± 495.06 non-germinated, and 255.23 ± 241.77 sterile pollen grains per pollen donor.

Mature fruits were collected five to six weeks after hand-pollination, yielding 112 fruits and 2,082 seeds. In the complementary experiment, seeds were classified as 85.5% viable, 1.0% germinated, 1.1% aborted, and 11.2% unfertilized. From each fruit, 40% of the viable seeds were randomly selected for sowing, with a minimum of five seeds per fruit, resulting in a total of 939 seeds. Seeds were sown individually into genotyping plate wells containing agar (10 g L⁻¹) to eliminate seed-density effects on germination, and yielded a germination rate of 99.4%. During paternity analyses, 54 offspring with missing data for three or more microsatellite markers were excluded. Paternity analyses were performed separately for each mating group, except for two groups in which two pollen donors were accidentally exchanged between the first and second pollination events. For these two groups, paternity analyses were performed separately for each pollination session, resulting in a total of eight independent paternity analyses at the scale of the complementary experiment. One of the eight microsatellite markers used for genotyping (m396) exhibited a high mismatch rate. Because mismatches were largely restricted to this marker and were not observed in the main experiment, they were likely due to capillary misalignment that emerged during the complementary experiment. Consequently, this marker was excluded from paternity analyses. Across the remaining seven markers, the mean mismatch rate was 0.098%, and the mean non- exclusion probability was 0.101, ranging from 0.063 to 0.168 across markers. Replacing brushes with toothpicks and systematically changing them between pollinations, substantially reduced unintended paternity attributed to pollen recipients to an average of 15.9%, ranging from 0.0% to 17.9% across the eight analyses. Paternity assignments attributed to recipient plants were excluded from statistical analyses, corresponding to 62 offspring. Of the 933 offspring genotyped, 704 paternity assignments were retained for statistical analyses.

### Statistical Analysis

Accounting for experimental variability in measured pollen and pistil traits

We first analyzed pollen and pistil traits independently using models that accounted for all known sources of experimental heterogeneity, in order to obtain trait proxies corrected for experimental artifacts. To this end, we constructed generalized linear mixed-effects models (GLMMs) using the spaMM package [75] (version 4.6.39) in R [76] (version 4.4.2), treating each measured trait as a response variable and including the structure of covariates and random factors described below. All analyses were carried out separately for the two experiments. Our models included fixed factors and uncontrolled experimental variables treated as random effects to account for potential sources of variation in pollen and pistil measurements. Predicted pollen and pistil traits values from these models were then used in subsequent analyses to (i) test for cryptic female choice and (ii) estimate the repeatability of pollen and pistil traits while statistically accounting for uncontrolled experimental variability. Hereafter, we refer to these predicted values as “proxies for pollen and pistil traits”.

Pollen density and image acquisition time were included as covariates in pollen trait models, as both factors are known to influence pollen growth dynamics [35,68,69]. Because pollen density present on microscope slides (*i.e.* the number of pollen grains counted per image) is often reported to have non-linear effects on pollen growth dynamics [68], we compared models including pollen density as either a linear or quadratic fixed effect using likelihood ratio tests (LRTs). Based on these comparisons, pollen density was included as a quadratic term for germination rate in both experiments, and for pollen size in the main experiment and pollen tube growth rate in the complementary experiment. Pollen trait models included pollen donor identity, sowing block, and stamen sampling date as random effects. In the complementary experiment, stamen identity was nested within flower identity, which was itself nested within donor plant identity. Pistil trait models included sowing block and pistil sampling date as random effects. In the main experiment, additional random effects were included: microscope objective and the source population of pollen donor plants in pollen trait models, and microscope type (binocular or optical microscope) and the source population of the pollen recipient in pistil traits model. Proxies for individual-level effects on pollen and pistil traits, which accounted for experimental artifacts, were obtained by extracting random effects from donor or recipient plant identity from GLMMs described above. We used such random effects as trait proxies because they capture among-individual variation while explicitly accounting for hierarchical structure and experimental heterogeneity, thereby providing unbiased individual-level estimates that are not confounded by other effects. These random effects were subsequently standardized prior to further analyses.

Pollen production was modeled as the number of pollen grains counted per collected stamen, assuming a Poisson-distributed residual variation. Pollen size and square-root-transformed pollen tube growth rate were analyzed using Gaussian error distributions. Germination rate was modeled using a binomial error distribution with a two-column response variable comprising (1) the number of germinated pollen grains and (2) the number of non-germinated pollen grains, including sterile pollen. Model predictions obtained with or without including sterile pollen grains were highly and positively correlated in both experiments (main experiment: r = 0.97, p < 0.001; complementary experiment: r = 0.99, p < 0.001), and subsequent analyses yielded consistent results regardless of whether sterile pollen was included.

Testing cryptic female choice through multinomial models of paternal fertilization succes In experimental designs where multiple sexual partners compete for access to ovules, paternity data are typically polytomous, and multinomial logit models are commonly used [77]. A key statistical challenge lies in accounting for correlations among paternity outcomes attributed to the same competitor, which arise because the fertilization success of a given pollen donor is necessarily dependent on the success of other donors competing for the same set of ovules. Data were analyzed using logit mixed models via a dedicated statistical procedure we developed, which is implemented in the novel pois4mlogit function available in the spaMM package [75]. This approach explicitly accounts for competitive dependencies among pollen donors within the same mating group, whose relative competitiveness varies because pollen donors were not selected based on prior knowledge on their fertilization ability. In our analysis, the multinomial sample size corresponded to the number of genotyped offspring per fruit from a given pollen recipient plant within a mating group.

Because the total number of pollen recipient and donor plants involved in hand-pollinations differed between the main and complementary experiments (70 versus 30), as did the offspring genotyping effort (100% versus 40%), and because minor protocol differences implied different random-effects structures, analyses were conducted separately for the two datasets using the same methodology. Prior to multinomial model construction, correlations among pollen and among pistil traits were assessed to identify independent phenotypic dimensions and minimize multicollinearity. Pairwise correlations were estimated using Spearman’s rank correlations on standardized RANEF-based trait proxies, and resulting *p*- values were adjusted for multiple testing using the FDR method [78]. Proxies of pollen size, germination rate, and tube growth rate were uncorrelated in both experiments, except for a positive but moderate correlation between pollen germination rate and pollen tube growth rate in the main experiment (Spearman’s ρ = 0.55, p < 0.001; Fig S2A and S2B). Many correlations between pistil traits were however found in both experiments but stigma area, style length, and ovary length were identified as three largely independent pistil traits, except for a positive correlation between ovary length and stigma area in the complementary experiment (Spearman’s ρ = 0.48, p = 0.021; Fig S1A and S1B). These traits represent distinct phenotypic dimensions of the pistil, capturing two independent measures of length and one measure of surface area, for which we had strong *a priori* expectations and were therefore retained for subsequent analyses. Longer pistil structures are expected to increase the distance pollen tubes must travel, whereas larger stigmatic surfaces are expected to increase pollen competition, potentially favoring faster-growing or more competitive pollen traits [6].

We first built multinomial models explaining variation in paternal fertilization success by including all three pollen trait proxies (standardized RANEF-based proxies for pollen size, pollen germination rate, and pollen tube growth rate) as fixed effects. All models included the covariate and random-effects structure described below. The effects of pollen trait proxies were tested by comparing models with and without the tested effect using likelihood ratio tests (LRTs). We then built models including interactions between pollen and pistil trait proxies using two complementary approaches: (i) “single-interaction” models including a single pollen trait and its interaction with a given pistil trait, and (ii) a complete model including all pollen traits and their interactions with the three retained pistil traits: stigma area, style length, and ovary length. Comparing these two types of models allowed us to assess the robustness of results and the influence of multicollinearity. Interaction effects were tested using LRTs, with p-values corrected for multiple testing using the FDR method [78] (*n* = 21 tests for single-interaction models; *n* = 9 tests for complete model). Pollen-by-pistil interactions obtained from the complete model are reported in the main text for the main experiment (Table 1) and in the Supporting Information for the complementary experiment (Table S6); all tested pollen-pistil interactions from single-interaction models in both experiments are reported in the Supporting Information (see Tables S5 and S7). We assessed the significance of pollen-pistil interactions across both experiments using Fisher’s combined probability test, applied separately to each model type (single-interaction and full models). This method combines p-values obtained independently that share the same null hypothesis – in our case testing whether a given pollen × pistil trait interaction is significant or not across both experiments [79]. A significant combined p-value therefore indicates a global effect of a given pollen-pistil interaction on paternal fertilization success across both experiments (Tables S8 and S9).

Each model included the relative pollen production of each competitor in the pollen mix used for manual pollinations as a covariate. Relative pollen production was defined as the pollen quantity produced by a given competitor (RANEF-based proxies) divided by the sum of pollen production across all competitors within the same mating group. In four mating groups of the main experiment, pollen production for one of the five pollen donors could not be estimated because dried pollen pellets could not be dissolved again (see above). In these cases, relative pollen production was calculated based on the four pollen donors for which pollen production could be reliably estimated, and multinomial logit models were applied to these four donors. The effect of pollen production on paternal fertilization success was tested using LRTs in models including germination rate, pollen size, and tube growth rate as main effects (Tables S1 and S2). In both experiments, pollen donor identity was included as a random effect to account for non-independence in fertilization success among fruits produced by the same pollen recipient. For the main experiment, the random-effects structure additionally included the source population of pollen donors in interaction with the source population of pollen recipient to account for potential effects of male-female coevolution on fertilization success.

Robustness analysis excluding potentially incompatible sexual partners To ensure that our results reflected interactions between compatible sexual partners, we conducted a sensitivity analysis to test the robustness of our findings to the exclusion of pollen donor that sired either 0%, ≤1%, or ≤5% of assigned paternity on a given pollen recipient. The rationale for excluding pollen donors with very low fertilization success is that such cases may reflect genetic incompatibility between donor and recipient rather than poor competitive ability. Although some level of selfing has been reported in the study line [56] – making it conceivable that genetically incompatible donor-recipient pairs occasionally produce limited seed set – this approach is highly conservative, as low fertilization success is more likely to reflect weak pollen competitive ability within mating groups. Exclusion thresholds were based on the proportion of seeds sired by each pollen donor across all fruits collected from a given pollen recipient plant (main experiment: *n* = 2 fruits; complementary experiment: *n* = 4 fruits). Because results were similar across the three exclusion thresholds for the main experiment, and consistent across thresholds for the complementary experiment, we report only the results obtained using the ≤5% threshold in the Supporting Information (see Tables S10–S13).

Repeatability of pollen and pistil traits

Repeatability of pollen and pistil traits was estimated using LMMs implemented in the rptR package [80] (version 0.9.2) in R, using as response values for the repeatability models measures of variation of pollen and pistil traits, designed to tell apart individual-level variation from with-individual variation from other sources of experimental variability. Repeatability is defined as the proportion of total phenotypic variation attributable to differences among individuals relative to within-individual variation [81]. It is estimated as the proportion of variance explained by pollen donor or recipient plant identity relative to the total variance estimated by a linear mixed model with Gaussian residual variation.

Analyses were based on data collected from the complementary experiment, which provided extensive replication of pollen and pistil trait measurements across multiple flowers per pollen donor and recipient plant. For pistil traits, the response values of the repeatability models were the residuals of mixed-effect models for pistil measurements only including a fixed-effect intercept and random effects for experimental block and date. For pollen traits, the response values were, for each stamen, the sum of values of random effects at the individual pollen donor, flower, and stamen level, as predicted from a mixed effect model adjusted to pollen measurements from eight stamens collected from four flowers per pollen donor (*n* = 240 stamens) from the complementary experiment. The latter model also included random effects for experimental block, date, and experimenter. We report repeatability estimates with standard errors and bootstrapped 95% confidence intervals, based on 1,000 parametric bootstrap replicates implemented in the rpt function [80]. Parametric bootstrapping quantifies the uncertainty of repeatability estimates by simulating response variable from the fitted model followed by a re-estimation of repeatability.

## Acknowledgments

We are grateful to Denis Orcel for plant care, and to Elsa Noël and Julia Centanni for their work on pollen and pistil trait measurements. We are thankful to Fleur Hamoir for genotyping support. We also acknowledge the “PollenExtra” platform at ISEM (Montpellier, France) for providing access to microscopy equipment, and the “Plateforme des Terrains d’Expériences du LabEx CeMEB” (Montpellier, France) for logistical and technical support, including hosting plants during the experiment. Artificial intelligence tools were used solely to assist with rephrasing and improving the clarity of specific sentences.

## Author contributions

Conceptualization: JT Methodology: EB, TC, FR, JT, AM Software : FR

Formal Analysis : TC, FR Investigation: TC, EB, AM, JT Visualization: TC

Funding acquisition: JT

Project administration: JT, EB, TC Supervision: JT

Writing – original draft: TC, JT

Writing – review & editing: TC, JT, FR, EB, AM

## Data, code, and materials availability

The original datasets, scripts, and all materials used to generate the results and figures are publicly available at Zenodo: https://doi.org/10.5281/zenodo.20545213

