## Supporting Information for "Experimental evidence for female choice in an angiosperm"

**S1 Text. Multinomial procedure for analysing pollen competition experiments.** Details of multinomial models.

In experimental designs where more than two sexual partners compete for access to ovules, paternity data are typically polytomous. Here the observations are counts $n_{ic}$ of the number of paternities from different pollen donors ($c = 1, \ldots, C$) in different multinomial draws ($i = 1, \ldots, n$ for distinct fruits). The multinomial logit model can be used to fit such data, as in analyses of other mate choice experiments [1,2]. In such a model, the expected frequencies $p_{ic}$ of the different types $c = 1, \ldots, C$ in different multinomial draws ($i = 1, \ldots, n$ for, say, different focal mother plants or even distinct fruits) are written

$$p_{ic} = \frac{e^{\eta_{ic}}}{\sum_{c=1}^{C} e^{\eta_{ic}}}$$

where each $\exp(\eta_{ic})$ quantifies the relative ability of each competitor to be the father. Each $\eta_{ic}$ takes the form of a linear predictor as considered in linear, generalized linear, and generalized linear mixed models (GLMMs). In the GLMM case in particular, it is a sum of fixed and random effects. A procedure, `pois4mlogit`, has been implemented in the `spaMM` R package for mixed-effect models [3], allowing to fit and compare multinomial logit models, including those with the additional features described below. This procedure is based on the concept of fitting a surrogate multivariate-response Poisson GLM or GLMM corrected so that the sums, over competing partners for each multinomial draw, of the expected values from the adjusted Poisson distributions, equal the multinomial sample sizes [4]. Further details on the fitting strategy can be found in the documentation of the `pois4mlogit` function.

In the present analyses, the competing pollen donors are exchangeable: the likelihood should be the same whatever the indexing (ordering) of these donors within each multinomial draw. This implies a number of constraints that should be respected by the fitting procedure. For example, any predictor variable (such as the pollen tube growth rate) should have the same fitted coefficient in the linear predictors for the different types (*i.e.*, the different pollen donors). Likewise, any random effect intrinsic to a competing partner (*e.g.,* a genetic effect of the pollen donor) included in a linear predictor will appear identically in all linear predictors for the five competing partners and should have a single fitted value of its variance, rather than five distinct fitted variances. Further, we need to be able to specify that any random effect intrinsic to the pollen donor (such as its genetic effect) takes the same value in the different contexts of interaction of this donor, for example when it is the “first” partner in interaction with a given mother plant and the “third” partner in interaction with another mother plant.

These constraints also have further implications for the terms that can usefully be included in the linear predictors. In particular, if one includes an intercept term in the linear predictor, it should be identical for the different partners, and then the form of $p_{ic}$ in the above equation implies that the intercept has no effect on the $p_{ic}$’s nor on the likelihood function. Likewise, a mother plant’s trait should act identically on the success of all competing partners in a competition arena, and thus it will not affect the likelihood, unless this trait acts in interaction with a partner’s trait (with a distinct value for each pollen donor). Thus, the linear predictors can include an interaction term with the mother plant’s trait even though they do not involve the mother plant’s trait in isolation as one of the terms of each linear predictor.

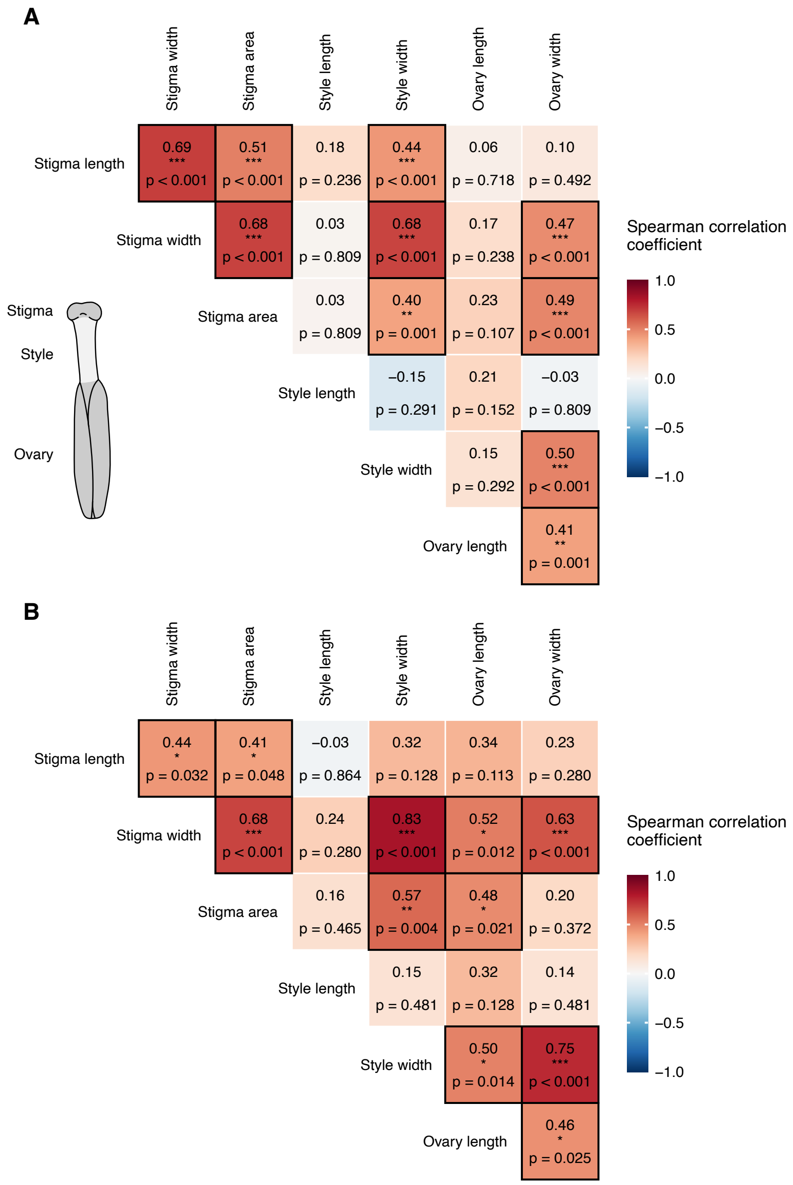


**Fig S1. Heatmap of correlations among pistil traits.** Pistil traits were standardized and correspond to predicted effects extracted from separate GLMMs correcting for potentially confounding experimental factors treated as random effects. Panels show pairwise correlations among pistil traits estimated using Spearman's rank correlation for **(A)** the main experiment, with extensive replication of parental plants, and **(B)** the complementary experiment, with extensive replication of pistil trait measurements per pollen recipient. Color intensity reflects the strength and direction of correlations. *P*-values were adjusted for multiple testing using the FDR method [5] ($n = 21$ tests). Significant correlations are outlined in black. Asterisks represent significance levels: $P < 0.05$ (*), $P < 0.01$ (**), $P < 0.001$ (***).

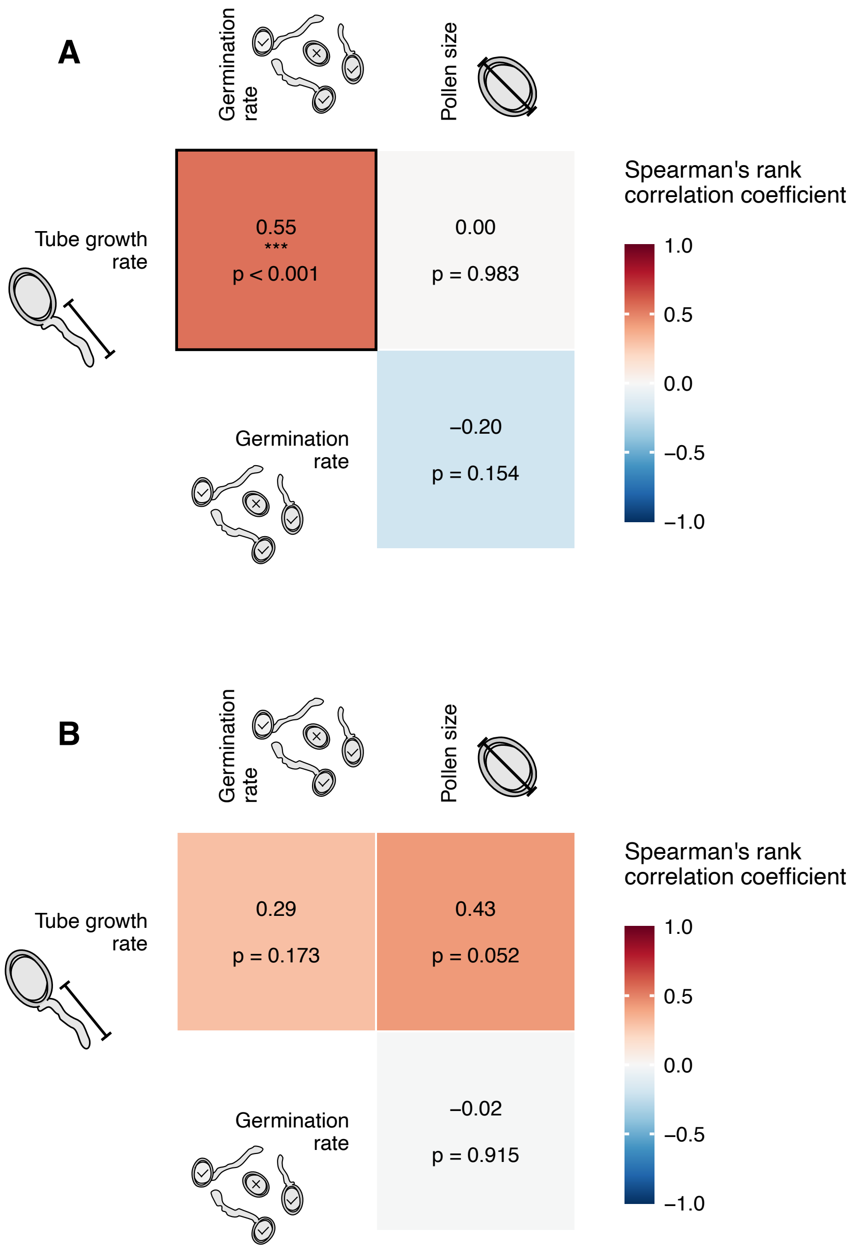


**Fig S2. Heatmap of correlations among pollen traits.** Pollen traits were standardized and represent predicted effects extracted from separate GLMMs correcting for potentially confounding experimental factors treated as random effects. Panels show pairwise correlations among pollen traits estimated using Spearman's rank correlation for **(A)** the main experiment with extensive replication of parental plants, and **(B)** the complementary experiment with extensive replication of pollen trait measurements across multiple flowers per pollen donor. Color intensity reflects the strength and direction of correlations. *P*-values were adjusted for multiple testing using the FDR method [5] ($n = 3$ tests). Significant correlations are outlined in black. Asterisks represent significance levels: $P < 0.05$ (*), $P < 0.01$ (**), $P < 0.001$ (***).

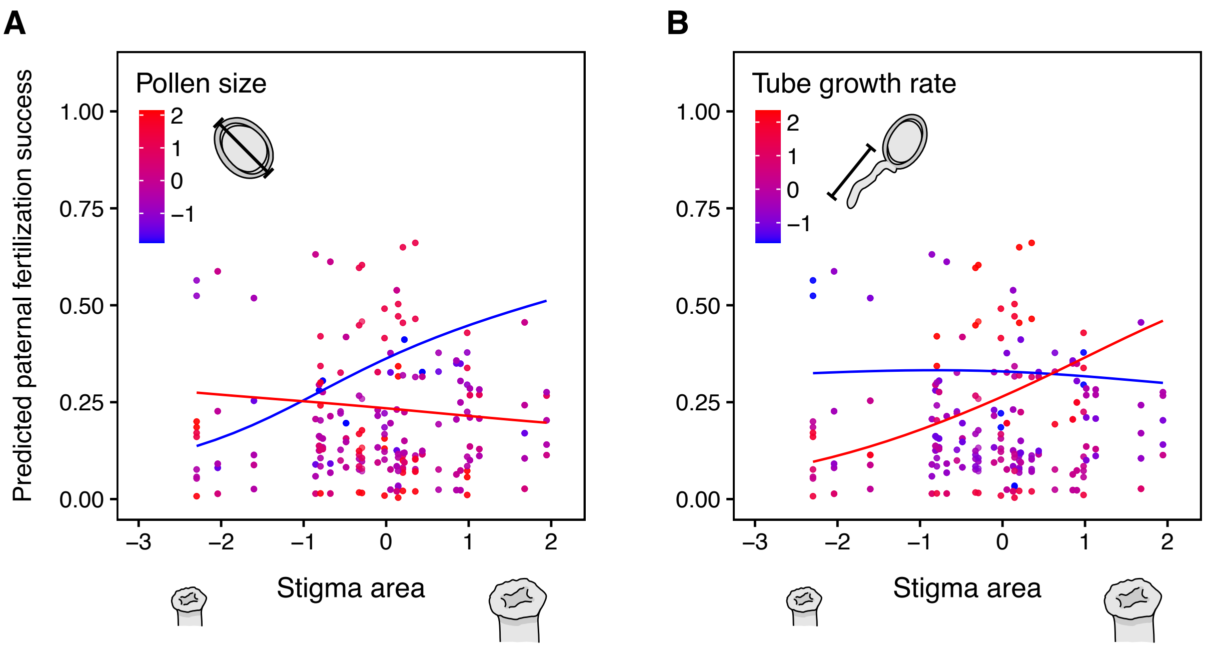


**Fig S3. Opposite-directional pollen-pistil trait interactions on predicted paternal fertilization success.** Points show predicted fertilization success of pollen donor plants across pistil trait values, with pollen trait values indicated by a blue-to-red gradient. Blue and red lines indicate model predictions for low (0.1 quantile) and high (0.9 quantile) pollen trait values, respectively. Panels display significant interactions between **(A)** pollen size and stigma area and **(B)** tube growth rate and stigma area. Analyses were based on the complementary experiment, which included extensive replication of pollen measurements per pollen donor. The model included germination rate, pollen size, and tube growth rate as main effects, and their interaction with stigma area, style length and ovary length. Relative pollen production was included as a covariate, and pollen donor plant identity and source population were included as random effects. All pollen and pistil traits were standardized and correspond to predicted values extracted from individual-level random effects of GLMMs that corrected for pollen density and other potentially confounding experimental factors included as random effects.

**Table S1. Estimated directional effects of pollen traits on paternal fertilization success for the main experiment.**

| Pollen trait | $\beta \pm$ SE | $\chi^2$ | df | p-value (LRT) |
|---|---|---|---|---|
| Germination rate | 0.354 ± 0.197 | 3.101 | 1 | 0.078 |
| Pollen size | −0.166 ± 0.155 | 1.116 | 1 | 0.291 |
| Tube growth rate | −0.128 ± 0.184 | 0.473 | 1 | 0.492 |
| Pollen production | 1.023 ± 1.059 | 0.904 | 1 | 0.342 |

Each pollen trait main effect was tested using LRTs comparing models with and without the focal pollen trait term. Analyses were based on the main experiment with extensive replication of parental plants. The model included germination rate, pollen size, tube growth rate, and the relative pollen production of competitors as a covariate, with pollen donor plant identity and source population included as random effects. All pollen traits were standardized and correspond to predicted values extracted from individual-level random effects of generalized linear mixed-effects models (GLMMs) that corrected for pollen density and other potentially confounding experimental factors included as random effects.

**Table S2. Estimated directional effects of traits on paternal fertilization success for the complementary experiment.**

| Pollen trait | β ± SE | $\chi^2$ | df | p-value (LRT) |
|---|---|---|---|---|
| Germination rate | 0.366 ± 0.229 | 2.487 | 1 | 0.115 |
| Pollen size | −0.240 ± 0.251 | 0.909 | 1 | 0.340 |
| Tube growth rate | −0.130 ± 0.253 | 0.263 | 1 | 0.608 |
| Pollen production | 1.089 ± 2.756 | 0.153 | 1 | 0.696 |

Each pollen trait main effect was tested using LRTs comparing models with and without the focal pollen trait term. Analyses were based on the complementary experiment with extensive replication of pollen and pistil trait measurements across multiple flowers per parental plant. The model included germination rate, pollen size, tube growth rate, and the relative pollen production of competitors as a covariate, with pollen donor plant identity and source population included as random effects. All pollen traits were standardized and correspond to predicted values extracted from individual-level random effects of GLMMs that corrected for pollen density and other potentially confounding experimental factors included as random effects.

**Table S3. Estimated linear and quadratic effects of pollen density on pollen trait variation from the main experiment.**

| Pollen density effect | β | SE | $\chi^2$ | df | p-value (LRT) |
|---|---|---|---|---|---|
| **Germination rate** | | | | | |
| Linear | $-1.894 \times 10^{-3}$ | $1.458 \times 10^{-4}$ | 2.604 | 1 | < 0.001* |
| Quadratic | $-1.656 \times 10^{-6}$ | $3.194 \times 10^{-7}$ | 2.794 | 1 | < 0.001* |
| **Pollen size (μm)** | | | | | |
| Linear | $1.82 \times 10^{-3}$ | $3.226 \times 10^{-3}$ | 3.100 | 1 | 0.577 |
| Quadratic | $-8.995 \times 10^{-5}$ | $4.840 \times 10^{-5}$ | 3.400 | 1 | 0.065 |
| **Tube growth rate (μm)** | | | | | |
| Linear | $-3.646 \times 10^{-3}$ | $2.45 \times 10^{-3}$ | 2.190 | 1 | 0.139 |
| Quadratic | $-5.862 \times 10^{-5}$ | $3.644 \times 10^{-5}$ | 2.584 | 1 | 0.108 |

Linear and quadratic effects of pollen density were tested separately for each pollen trait (germination rate, pollen size, and pollen tube growth rate) using LRTs comparing models with and without the effect. Analyses were based on the main experiment, which included extensive replication of parental plants. Linear effects were tested in GLMMs excluding the quadratic term, whereas quadratic effects were tested in models including the linear term. Models accounted for repeated measurements over time as a covariate and included potentially confounding experimental factors as random effects. Gaussian error distributions were assumed for pollen size and square-root-transformed tube growth rate, and a binomial error distribution was used for germination rate. Asterisks represent statistically significant *p*-values.

**Table S4. Estimated linear and quadratic effects of pollen density on pollen trait variation from the complementary experiment.**

| Pollen density effect | β | SE | $\chi^2$ | df | p-value (LRT) |
|---|---|---|---|---|---|
| **Germination rate** | | | | | |
| Linear | $7.781 \times 10^{-4}$ | $9.611 \times 10^{-5}$ | 6.556 | 1 | < 0.001* |
| Quadratic | $1.053 \times 10^{-6}$ | $4.491 \times 10^{-7}$ | 5.475 | 1 | 0.019* |
| **Pollen size (µm)** | | | | | |
| Linear | $4.271 \times 10^{-4}$ | $3.267 \times 10^{-5}$ | 1.685 | 1 | 0.194 |
| Quadratic | $-5.163 \times 10^{-7}$ | $1.772 \times 10^{-6}$ | 8.500 | 1 | 0.771 |
| **Tube growth rate (µm)** | | | | | |
| Linear | $-4.957 \times 10^{-4}$ | $2.966 \times 10^{-4}$ | 2.770 | 1 | 0.096 |
| Quadratic | $4.995 \times 10^{-6}$ | $1.473 \times 10^{-6}$ | 1.146 | 1 | < 0.001* |

Linear and quadratic effects of pollen density were tested separately for each pollen trait (germination rate, pollen size, and pollen tube growth rate) using LRTs comparing models with and without the effect. Analyses were based on the complementary experiment, which included extensive replication of pollen measurements per pollen donor. Linear effects were tested in GLMMs excluding the quadratic term, whereas quadratic effects were tested in models including the linear term. Models accounted for repeated measurements over time as a covariate and included potentially confounding experimental factors as random effects. Gaussian error distributions were assumed for pollen size and square-root-transformed tube growth rate, and a binomial error distribution was used for germination rate. Asterisks represent statistically significant *p*-values.

**Table S5. Estimated effects of pollen by pistil trait interactions on paternal fertilization success in single-interaction models from the complementary experiment.**

| Pollen trait × Pistil trait | β ± SE | $\chi^2$ | df | p-value (LRT) | FDR p-value |
|---|---|---|---|---|---|
| **Germination rate** | | | | | |
| Germination rate × Stigma length | 0.112 ± 0.052 | 4.497 | 1 | 0.034* | 0.055 |
| Germination rate × Stigma width | 0.177 ± 0.046 | 15.317 | 1 | < 0.001* | < 0.001* |
| Germination rate × Stigma area | 0.151 ± 0.044 | 12.155 | 1 | < 0.001* | 0.002* |
| Germination rate × Style length | −0.184 ± 0.052 | 11.962 | 1 | < 0.001* | 0.002* ‡ |
| Germination rate × Style width | 0.112 ± 0.043 | 6.554 | 1 | 0.010* | 0.024* |
| Germination rate × Ovary length | 0.147 ± 0.056 | 7.089 | 1 | 0.008* | 0.020* † |
| Germination rate × Ovary width | 0.206 ± 0.050 | 17.389 | 1 | < 0.001* | 0.001* † |
| **Pollen size** | | | | | |
| Pollen size × Stigma length | −0.015 ± 0.065 | 0.055 | 1 | 0.815 | 0.815 |
| Pollen size × Stigma width | 0.022 ± 0.066 | 0.111 | 1 | 0.739 | 0.815 |
| Pollen size × Stigma area | −0.095 ± 0.056 | 2.820 | 1 | 0.093 | 0.140 |
| Pollen size × Style length | −0.117 ± 0.055 | 4.518 | 1 | 0.034* | 0.055 |
| Pollen size × Style width | 0.182 ± 0.068 | 7.159 | 1 | 0.007* | 0.020* |
| Pollen size × Ovary length | 0.019 ± 0.065 | 0.080 | 1 | 0.778 | 0.815 |
| Pollen size × Ovary width | −0.076 ± 0.063 | 1.405 | 1 | 0.236 | 0.300 |
| **Tube growth rate** | | | | | |
| Tube growth rate × Stigma length | 0.131 ± 0.061 | 4.502 | 1 | 0.034* | 0.055 |
| Tube growth rate × Stigma width | 0.202 ± 0.055 | 13.342 | 1 | < 0.001* | 0.002* |
| Tube growth rate × Stigma area | 0.082 ± 0.053 | 2.357 | 1 | 0.125 | 0.175 |
| Tube growth rate × Style length | −0.049 ± 0.045 | 1.144 | 1 | 0.285 | 0.332 |
| Tube growth rate × Style width | 0.103 ± 0.042 | 5.806 | 1 | 0.016* | 0.034* |
| Tube growth rate × Ovary length | 0.055 ± 0.047 | 1.364 | 1 | 0.243 | 0.300 |
| Tube growth rate × Ovary width | 0.138 ± 0.045 | 9.199 | 1 | 0.002* | 0.008* |

Each pollen-pistil trait interaction was tested separately using LRTs comparing models with and without the interaction term. Analyses were based on the complementary experiment, which included extensive replication of pollen and pistil trait measurements across multiple flowers per parental plant. In addition to the pollen-pistil interaction term, single-interaction models included the focal pollen trait, the relative pollen production of competitors as a covariate, and pollen donor identity as a random effect. *P*-values were adjusted for multiple testing across all interactions ($n$ = 21) using the FDR method [5]. All pollen and pistil traits were standardized and correspond to predicted values extracted from individual-level random effects of GLMMs that corrected for pollen density and other potentially confounding experimental factors included as random effects. Asterisks represent significant *p*-values. The † symbol indicates significant pollen-pistil interactions that are consistent across the two experiments comparing models with identical fixed-effect structure, whereas the ‡ symbol indicates interactions with opposite significant effects (see Table S5).

**Table S6. Estimated effects of pollen by pistil traits interaction on paternal fertilization success in the complete model from the complementary experiment.**

| Pollen trait × Pistil trait | β ± SE | $\chi^2$ | df | p-value (LRT) | FDR p-value |
|---|---|---|---|---|---|
| **Germination rate** | | | | | |
| Germination rate × Stigma area | −0.186 ± 0.089 | 4.347 | 1 | 0.037* | 0.083 |
| Germination rate × Style length | 0.076 ± 0.090 | 0.718 | 1 | 0.397 | 0.446 |
| Germination rate × Ovary length | 0.236 ± 0.093 | 6.538 | 1 | 0.011* | 0.032* † |
| **Pollen size** | | | | | |
| Pollen size × Stigma area | −0.288 ± 0.093 | 9.602 | 1 | 0.002* | 0.009* |
| Pollen size × Style length | 0.102 ± 0.084 | 1.485 | 1 | 0.223 | 0.287 |
| Pollen size × Ovary length | 0.148 ± 0.098 | 2.269 | 1 | 0.132 | 0.198 |
| **Tube growth rate** | | | | | |
| Tube growth rate × Stigma area | 0.258 ± 0.079 | 10.856 | 1 | 0.001* | 0.009* |
| Tube growth rate × Style length | −0.102 ± 0.066 | 2.346 | 1 | 0.126 | 0.198 |
| Tube growth rate × Ovary length | −0.038 ± 0.075 | 0.252 | 1 | 0.616 | 0.616 |

Each pollen-pistil trait interaction was tested using LRTs comparing models with and without the interaction term. Analyses were based on the complementary experiment, which included extensive replication of pollen and pistil trait measurements across multiple flowers per parental plant. The model included germination rate, pollen size or tube growth rate as main effects, along with their interaction with stigma area, style length and ovary length. Relative pollen production of competitors was included as a covariate, and pollen donor plant identity and source population were included as random effects. *P*-values were adjusted for multiple testing across all interactions ($n = 9$) using the FDR method [5]. Only interactions involving uncorrelated components of pistil morphology are reported in the main text (stigma area, style length, and ovary length; Fig. S1; Table S7). All pollen and pistil traits were standardized and correspond to predicted values extracted from individual-level random effects of GLMMs that corrected for pollen density and other potentially confounding experimental factors included as random effects. Asterisks represent significant *p*-values. The † symbol indicates significant pollen-pistil interactions that are consistent across the two experiments comparing models with identical fixed-effect structure (see Table 1).

**Table S7. Estimated effects of pollen by pistil trait interactions on paternal fertilization success in single-interaction models from the complementary experiment.**

| Pollen trait × Pistil trait | β ± SE | $\chi^2$ | df | p-value (LRT) | FDR p-value |
|---|---|---|---|---|---|
| **Germination rate** | | | | | |
| Germination rate × Stigma length | −0.012 ± 0.037 | 0.097 | 1 | 0.755 | 0.755 |
| Germination rate × Stigma width | 0.083 ± 0.047 | 3.082 | 1 | 0.079 | 0.151 |
| Germination rate × Stigma area | 0.074 ± 0.048 | 2.387 | 1 | 0.122 | 0.184 |
| Germination rate × Style length | 0.131 ± 0.056 | 6.174 | 1 | 0.013* | 0.034* ‡ |
| Germination rate × Style width | 0.043 ± 0.039 | 1.211 | 1 | 0.271 | 0.335 |
| Germination rate × Ovary length | 0.152 ± 0.042 | 13.540 | 1 | < 0.001* | 0.002* † |
| Germination rate × Ovary width | 0.131 ± 0.046 | 7.928 | 1 | 0.005* | 0.017* † |
| **Pollen size** | | | | | |
| Pollen size × Stigma length | −0.015 ± 0.046 | 0.100 | 1 | 0.752 | 0.755 |
| Pollen size × Stigma width | 0.067 ± 0.058 | 1.339 | 1 | 0.247 | 0.324 |
| Pollen size × Stigma area | −0.029 ± 0.058 | 0.237 | 1 | 0.626 | 0.692 |
| Pollen size × Style length | 0.163 ± 0.058 | 8.281 | 1 | 0.004* | 0.017* |
| Pollen size × Style width | 0.026 ± 0.050 | 0.269 | 1 | 0.604 | 0.692 |
| Pollen size × Ovary length | 0.143 ± 0.053 | 7.477 | 1 | 0.006* | 0.019* |
| Pollen size × Ovary width | 0.085 ± 0.051 | 2.741 | 1 | 0.098 | 0.171 |
| **Tube growth rate** | | | | | |
| Tube growth rate × Stigma length | 0.175 ± 0.042 | 18.920 | 1 | < 0.001* | < 0.001* |
| Tube growth rate × Stigma width | 0.097 ± 0.042 | 5.246 | 1 | 0.022* | 0.051 |
| Tube growth rate × Stigma area | 0.161 ± 0.050 | 10.955 | 1 | 0.001* | 0.005* |
| Tube growth rate × Style length | 0.074 ± 0.048 | 2.418 | 1 | 0.120 | 0.184 |
| Tube growth rate × Style width | 0.078 ± 0.036 | 4.649 | 1 | 0.031* | 0.065 |
| Tube growth rate × Ovary length | 0.160 ± 0.044 | 13.777 | 1 | 0.001* | 0.002* |
| Tube growth rate × Ovary width | −0.051 ± 0.039 | 1.742 | 1 | 0.187 | 0.262 |

Each pollen-pistil trait interaction was tested separately using LRTs comparing models with and without the interaction term. Analyses were based on the complementary experiment, which included extensive replication of pollen and pistil trait measurements across multiple flowers per parental plant. In addition to the pollen-pistil interaction term, single-interaction models included the focal pollen trait, the relative pollen production of competitors as a covariate, and pollen donor identity as a random effect. *P*-values were adjusted for multiple testing across all interactions ($n$ = 21) using the FDR method [5]. All pollen and pistil traits were standardized and correspond to predicted values extracted from individual-level random effects of GLMMs that corrected for pollen density and other potentially confounding experimental factors included as random effects. Asterisks represent significant *p*-values. The † symbol indicates significant pollen-pistil interactions that are consistent across the two experiments comparing models with identical fixed-effect structure, whereas the ‡ symbol indicates interactions with opposite significant effects (see Table S5).

**Table S8. Effect of pollen by pistil trait interactions on paternal fertilization success in the complete model from the main and the complementary experiments, with results obtained from Fisher's combined probability test.**

| | Main experiment | | Complementary experiment | | | |
|---|---|---|---|---|---|---|
| Pollen trait × Pistil trait | β ± SE | p-value (LRT) | β ± SE | p-value (LRT) | Fisher p-value | FDR Fisher p-value |
| **Germination rate** | | | | | | |
| Germination rate × Stigma area | 0.152 ± 0.059 | 0.008* | −0.186 ± 0.089 | 0.037* | 0.003* | 0.006* |
| Germination rate × Style length | −0.338 ± 0.082 | < 0.001* | 0.076 ± 0.090 | < 0.001* | < 0.001* | 0.001* |
| Germination rate × Ovary length | 0.290 ± 0.095 | 0.002* | 0.236 ± 0.093 | 0.011* | < 0.001 | < 0.001* |
| **Pollen size** | | | | | | |
| Pollen size × Stigma area | −0.021 ± 0.060 | 0.732 | −0.288 ± 0.093 | 0.002* | 0.011* | 0.014* |
| Pollen size × Style length | −0.195 ± 0.063 | 0.002* | 0.102 ± 0.084 | 0.223 | 0.003* | 0.006* |
| Pollen size × Ovary length | 0.109 ± 0.076 | 0.154 | 0.148 ± 0.098 | 0.132 | 0.099 | 0.112 |
| **Tube growth rate** | | | | | | |
| Tube growth rate × Stigma | −0.023 ± 0.071 | 0.744 | 0.258 ± 0.079 | 0.079 | 0.001* | 0.009* |
| Tube growth rate × Style length | 0.221 ± 0.072 | 0.002* | −0.102 ± 0.066 | 0.126 | 0.002* | 0.006* |
| Tube growth rate × Ovary length | −0.165 ± 0.083 | 0.046* | −0.038 ± 0.075 | 0.616 | 0.129 | 0.129 |

Each pollen-pistil trait interaction was tested separately using LRTs comparing the model with and without the interaction term for the main and the complementary experiments. The model included germination rate, pollen size or tube growth rate as main effects, along with their interaction with stigma area, style length and ovary length. Relative pollen production of competitors was included as a covariate, and pollen donor plant identity as a random effect. Source population was additionally included as a random effect in the complementary experiment. Only interactions involving uncorrelated components of pistil morphology are reported in the main text (stigma area, style length, and ovary length; Table S9). All pollen and pistil traits were standardized and correspond to predicted values extracted from individual-level random effects of generalized linear mixed-effects models (GLMMs) that corrected for pollen density and other potentially confounding experimental factors. Fisher's combined probability test [6] was applied to *p*-values obtained for each pollen-pistil interaction term across the two experiments. The resulting *p*-values were corrected using the FDR method [5]. Asterisks represent significant *p*-values.

**Table S9. Effect of pollen by pistil trait interactions on paternal fertilization success in single-interaction models from the main and the complementary experiments, with results obtained from Fisher's combined probability test.**

| | Main experiment | | Complementary experiment | | | |
|---|---|---|---|---|---|---|
| Pollen trait × Pistil trait | β ± SE | p-value (LRT) | β ± SE | p-value (LRT) | Fisher p-value | FDR Fisher p-value |
| **Germination rate** | | | | | | |
| Germination rate × Stigma length | 0.112 ± 0.052 | 0.034* | −0.012 ± 0.037 | 0.755 | 0.120 | 0.148 |
| Germination rate × Stigma width | 0.177 ± 0.046 | 0.001* | 0.083 ± 0.047 | 0.079 | < 0.001* | < 0.001* |
| Germination rate × Stigma area | 0.151 ± 0.044 | < 0.001* | 0.074 ± 0.048 | 0.122 | < 0.001* | 0.002* |
| Germination rate × Style length | −0.184 ± 0.052 | 0.001* | 0.131 ± 0.056 | 0.013* | 0.001* | 0.001* |
| Germination rate × Style width | 0.112 ± 0.043 | 0.010* | 0.043 ± 0.039 | 0.271 | 0.019* | 0.031* |
| Germination rate × Ovary length | 0.147 ± 0.056 | 0.008* | 0.152 ± 0.042 | < 0.001* | < 0.001* | < 0.001* |
| Germination rate × Ovary width | 0.206 ± 0.050 | 0.001* | 0.131 ± 0.046 | 0.005* | < 0.001* | < 0.001* |
| **Pollen size** | | | | | | |
| Pollen size × Stigma length | −0.015 ± 0.065 | 0.815 | −0.015 ± 0.046 | 0.752 | 0.913 | 0.913 |
| Pollen size × Stigma width | 0.022 ± 0.066 | 0.739 | 0.067 ± 0.058 | 0.247 | 0.493 | 0.518 |
| Pollen size × Stigma area | −0.095 ± 0.056 | 0.093 | −0.029 ± 0.058 | 0.626 | 0.224 | 0.248 |
| Pollen size × Style length | −0.117 ± 0.055 | 0.034* | 0.163 ± 0.058 | 0.004* | 0.001* | 0.003* |
| Pollen size × Style width | 0.182 ± 0.068 | 0.007* | 0.026 ± 0.050 | 0.604 | 0.029* | 0.043* |
| Pollen size × Ovary length | 0.019 ± 0.065 | 0.778 | 0.143 ± 0.053 | 0.006* | 0.031* | 0.043* |
| Pollen size × Ovary width | −0.076 ± 0.063 | 0.236 | 0.085 ± 0.051 | 0.098 | 0.110 | 0.144 |
| **Tube growth rate** | | | | | | |

| | | | | | | |
|---|---|---|---|---|---|---|
| Tube growth rate × Stigma length | 0.131 ± 0.061 | 0.034* | 0.175 ± 0.042 | < 0.001* | < 0.001* | < 0.001* |
| Tube growth rate × Stigma width | 0.202 ± 0.055 | < 0.001* | 0.097 ± 0.042 | 0.022* | < 0.001* | < 0.001* |
| Tube growth rate × Stigma area | 0.082 ± 0.053 | 0.125 | 0.161 ± 0.050 | < 0.001* | 0.001* | 0.003* |
| Tube growth rate × Style length | −0.049 ± 0.045 | 0.285 | 0.074 ± 0.048 | 0.120 | 0.149 | 0.174 |
| Tube growth rate × Style width | 0.103 ± 0.042 | 0.016* | 0.078 ± 0.036 | 0.031* | 0.004* | 0.007* |
| Tube growth rate × Ovary length | 0.055 ± 0.047 | 0.243 | 0.160 ± 0.044 | < 0.001* | < 0.001* | 0.002* |
| Tube growth rate × Ovary width | 0.138 ± 0.045 | 0.002* | −0.051 ± 0.039 | 0.187 | 0.004* | 0.007* |

Each pollen-pistil trait interaction was tested separately using LRTs comparing the model with and without the interaction term for the main and the complementary experiments. In addition to the pollen-pistil interaction term, single-interaction models included the focal pollen trait, the relative pollen production of competitors as a covariate, and pollen donor plant identity as a random effect. Source population was additionally included as a random effect in the complementary experiment. All pollen and pistil traits were standardized and correspond to predicted values extracted from individual-level random effects of generalized linear mixed-effects models (GLMMs) that corrected for pollen density and other potentially confounding experimental factors. Fisher's combined probability test [6] was applied to $p$-values obtained for each pollen-pistil interaction term across the two experiments. The resulting $p$-values were corrected using the FDR method [5]. Asterisks represent significant $p$-values.

**Table S10. Observed and estimated repeatabilities of pollen traits.**

| | | | Bootstrapping | |
|---|---|---|---|---|
| Pollen trait | Level | Observed RPT ± SE | Estimated RPT | 95% CI |
| Germination speed | Plant | 0.227 ± 0.082 | 0.222 | [0.056–0.375] |
| | Flower | 0.273 ± 0.084 | 0.273 | [0.114–0.433] |
| Pollen size | Plant | 0.691 ± 0.071 | 0.678 | [0.528–0.801] |
| | Flower | 0.271 ± 0.065 | 0.282 | [0.170–0.422] |
| Tube growth rate | Plant | 0.322 ± 0.087 | 0.319 | [0.153–0.478] |
| | Flower | 0.233 ± 0.077 | 0.229 | [0.090–0.393] |

Pollen trait repeatabilities at both the plant and flower levels were estimated using generalized linear mixed-effects models (GLMMs implemented via the `rpt` function in the `rptR` package [7]). Models treated pollen traits as response variables (standardized and extracted from GLMMs correcting for relevant confounding experimental covariates), with pollen donor plant identity and flower identity included as random factors. Gaussian error distributions were assumed for all pollen traits. Analyses were based on a complementary experiment, which featured extensive replication of pollen measurements across multiple flowers per pollen donor. Observed repeatabilities were derived directly from the variance components estimated by the models. Repeatability estimates correspond to mean values obtained from 1,000 parametric bootstrap simulations, with associated 95% confidence intervals, and are presented on the original scale.

**Table S11. Observed and estimated repeatabilities of pistil traits.**

| | | Bootstrapping | |
|---|---|---|---|
| Pistil trait | Observed RPT ± SE | Estimated RPT | 95% CI |
| Stigma length | 0.263 ± 0.094 | 0.259 | [0.080–0.443] |
| Stigma width | 0.474 ± 0.091 | 0.462 | [0.264–0.623] |
| Stigma area | 0.354 ± 0.095 | 0.346 | [0.154–0.527] |
| Style length | 0.426 ± 0.096 | 0.416 | [0.223–0.593] |
| Style width | 0.474 ± 0.093 | 0.470 | [0.268–0.629] |
| Ovary length | 0.368 ± 0.100 | 0.361 | [0.153–0.537] |
| Ovary width | 0.471 ± 0.096 | 0.459 | [0.244–0.627] |

Pistil trait repeatabilities were estimated using generalized linear mixed-effects models (GLMMs implemented via the `rpt` function in the `rptR` package [7]). Models treated pistil traits as response variables (standardized and extracted from GLMMs correcting for relevant confounding experimental covariates), with pollen recipient plant identity and flower identity included as random factors. Gaussian error distributions were assumed for all pollen traits. Analyses were based on a complementary experiment, which featured extensive replication of pistil measurements across multiple flowers per pollen recipient. Observed repeatabilities were derived directly from the variance components estimated by the models. Repeatability estimates correspond to mean values obtained from 1,000 parametric bootstrap simulations, with associated 95% confidence intervals, and are presented on the original scale.

**Table S12. Estimated effects of pollen by pistil trait interactions on paternal fertilization success in the complete model from the main experiment, excluding paternal plants with less than 5% of fertilized seeds.**

| Pollen trait × Pistil trait | β ± SE | $\chi^2$ | df | p-value (LRT) | FDR p-value |
|---|---|---|---|---|---|
| **Germination rate** | | | | | |
| Germination rate × Stigma area | 0.037 ± 0.059 | 0.386 | 1 | 0.534 | 0.801 |
| Germination rate × Style length | −0.177 ± 0.089 | 3.954 | 1 | 0.047* | 0.123 |
| Germination rate × Ovary length | 0.199 ± 0.103 | 3.696 | 1 | 0.055 | 0.123 |
| **Pollen size** | | | | | |
| Pollen size × Stigma area | 0.053 ± 0.062 | 0.677 | 1 | 0.411 | 0.739 |
| Pollen size × Style length | 0.005 ± 0.065 | 0.005 | 1 | 0.943 | 0.943 |
| Pollen size × Ovary length | −0.037 ± 0.081 | 0.200 | 1 | 0.655 | 0.842 |
| **Tube growth rate** | | | | | |
| Tube growth rate × Stigma area | 0.211 ± 0.075 | 7.468 | 1 | 0.006* | 0.035* |
| Tube growth rate × Style length | 0.013 ± 0.085 | 0.021 | 1 | 0.885 | 0.943 |
| Tube growth rate × Ovary length | −0.229 ± 0.086 | 7.084 | 1 | 0.008* | 0.035* |

Each pollen-pistil trait interaction was tested separately using LRTs comparing the model with and without the interaction term. Analyses were based on the main experiment with extensive replication of parental plants; pollen donor plants contributing less than 5% of fertilized seeds on a given pollen recipient plant were excluded from the analyses. The model included germination rate, pollen size or tube growth rate as main effects, along with their interaction with stigma area, style length and ovary length. Relative pollen production of competitors was included as a covariate, and pollen donor plant identity and source population were included as random effects. *P*-values were adjusted for multiple testing across all interactions ($n = 9$) using the FDR method [5]. Only interactions involving uncorrelated components of pistil morphology are reported in the main text (stigma area, style length, and ovary length; Table S5). All pollen and pistil traits were standardized and correspond to predicted values extracted from individual-level random effects of generalized linear mixed-effects models (GLMMs) that corrected for pollen density and other potentially confounding experimental. Asterisks represent significant *p*-values.

**Table S13. Estimated effects of pollen by pistil trait interactions on paternal fertilization success in single-interaction models from the main experiment, excluding paternal plants with less than 5% of fertilized seeds.**

| Pollen trait × Pistil trait | β ± SE | $\chi^2$ | df | p-value (LRT) | FDR p-value |
|---|---|---|---|---|---|
| **Germination rate** | | | | | |
| Germination rate × Stigma length | 0.033 ± 0.054 | 0.356 | 1 | 0.551 | 0.667 |
| Germination rate × Stigma width | 0.120 ± 0.048 | 6.685 | 1 | 0.010* | 0.049* |
| Germination rate × Stigma area | 0.107 ± 0.047 | 5.332 | 1 | 0.021* | 0.063 |
| Germination rate × Style length | −0.155 ± 0.052 | 8.958 | 1 | 0.003* | 0.029* |
| Germination rate × Style width | 0.081 ± 0.043 | 3.568 | 1 | 0.059 | 0.112 |
| Germination rate × Ovary length | 0.023 ± 0.061 | 0.134 | 1 | 0.714 | 0.789 |
| Germination rate × Ovary width | 0.108 ± 0.053 | 4.189 | 1 | 0.041* | 0.094 |
| **Pollen size** | | | | | |
| Pollen size × Stigma length | 0.090 ± 0.072 | 1.522 | 1 | 0.217 | 0.306 |
| Pollen size × Stigma width | 0.117 ± 0.078 | 2.095 | 1 | 0.148 | 0.239 |
| Pollen size × Stigma area | −0.014 ± 0.062 | 0.049 | 1 | 0.824 | 0.865 |
| Pollen size × Style length | 0.009 ± 0.056 | 0.026 | 1 | 0.872 | 0.872 |
| Pollen size × Style width | 0.114 ± 0.066 | 2.871 | 1 | 0.090 | 0.158 |
| Pollen size × Ovary length | −0.068 ± 0.077 | 0.765 | 1 | 0.382 | 0.501 |
| Pollen size × Ovary width | −0.040 ± 0.069 | 0.320 | 1 | 0.571 | 0.667 |
| **Tube growth rate** | | | | | |
| Tube growth rate × Stigma length | 0.128 ± 0.059 | 4.690 | 1 | 0.030* | 0.080 |
| Tube growth rate × Stigma width | 0.202 ± 0.060 | 11.644 | 1 | < 0.001* | 0.014* |
| Tube growth rate × Stigma area | 0.146 ± 0.060 | 5.861 | 1 | 0.015* | 0.054 |
| Tube growth rate × Style length | −0.096 ± 0.047 | 4.027 | 1 | 0.045* | 0.094 |
| Tube growth rate × Style width | 0.107 ± 0.042 | 6.418 | 1 | 0.011* | 0.049* |
| Tube growth rate × Ovary length | −0.063 ± 0.051 | 1.513 | 1 | 0.219 | 0.306 |

Each pollen-pistil trait interaction was tested separately using LRTs comparing models with and without the interaction term. Analyses were based on the main experiment with extensive replication of parental plants; pollen donor plants contributing less than 5% of fertilized seeds on a given pollen recipient plant were excluded from the analyses. In addition to the pollen-pistil interaction term, single-interaction models included the focal pollen trait, the relative pollen production of competitors as a covariate, and pollen donor plant identity as a random effect. *P*-values were adjusted for multiple testing across all interactions ($n = 21$) using the FDR method [5]. All pollen and pistil traits were standardized and correspond to predicted values extracted from individual-level random effects of GLMMs that corrected for pollen density and other potentially confounding experimental factors included as random effects. Asterisks represent significant *p*-values.

**Table S14. Estimated effects of pollen by pistil trait interactions on paternal fertilization success in the complete model from the complementary experiment, excluding paternal plants with less than 5% of fertilized seeds.**

| Pollen trait × Pistil trait | β ± SE | $\chi^2$ | df | p-value (LRT) | FDR p-value |
|---|---|---|---|---|---|
| **Germination rate** | | | | | |
| Germination rate × Stigma area | −0.031 ± 0.092 | 0.109 | 1 | 0.742 | 0.834 |
| Germination rate × Style length | −0.178 ± 0.113 | 2.441 | 1 | 0.118 | 0.266 |
| Germination rate × Ovary length | −0.006 ± 0.102 | 0.004 | 1 | 0.952 | 0.952 |
| **Pollen size** | | | | | |
| Pollen size × Stigma area | 0.095 ± 0.106 | 0.801 | 1 | 0.371 | 0.477 |
| Pollen size × Style length | 0.117 ± 0.110 | 1.109 | 1 | 0.292 | 0.439 |
| Pollen size × Ovary length | −0.158 ± 0.116 | 1.843 | 1 | 0.175 | 0.314 |
| **Tube growth rate** | | | | | |
| Tube growth rate × Stigma area | −0.184 ± 0.097 | 3.629 | 1 | 0.057 | 0.218 |
| Tube growth rate × Style length | −0.150 ± 0.083 | 3.225 | 1 | 0.073 | 0.218 |
| Tube growth rate × Ovary length | 0.287 ± 0.105 | 7.477 | 1 | 0.006* | 0.056 |

Each pollen-pistil trait interaction was tested separately using LRTs comparing models with and without the interaction term. Analyses were based on the complementary experiment, wich included extensive replication of pollen and pistil measurements per parental plant; pollen donor plants contributing less than 5% of fertilized seeds on a given pollen recipient plant were excluded from the analyses. The model included germination rate, pollen size or tube growth rate as main effects, along with their interaction with stigma area, style length and ovary length. Relative pollen production of competitors was included as a covariate, and pollen donor plant identity and source population were included as random effects. *P*-values were adjusted for multiple testing across all interactions ($n$ = 9) using the FDR method [5]. Only interactions involving uncorrelated components of pistil morphology are reported in the main text (stigma area, style length, and ovary length; Fig. S1). All pollen and pistil traits were standardized and correspond to predicted values extracted from individual-level random effects of GLMMs that corrected for pollen density and other potentially confounding experimental factors included as random effects. Asterisks represent significant *p*-values.

**Table S15. Estimated effects of pollen by pistil trait interactions on paternal fertilization success in single-interaction models from the complementary experiment, excluding paternal plants with less than 5% of fertilized seeds.**

| Pollen trait × Pistil trait | β ± SE | $\chi^2$ | df | p-value (LRT) | FDR p-value |
|---|---|---|---|---|---|
| **Germination rate** | | | | | |
| Germination rate × Stigma length | −0.044 ± 0.034 | 1.685 | 1 | 0.194 | 0.574 |
| Germination rate × Stigma width | −0.055 ± 0.044 | 1.494 | 1 | 0.222 | 0.574 |
| Germination rate × Stigma area | −0.018 ± 0.045 | 0.158 | 1 | 0.691 | 0.918 |
| Germination rate × Style length | −0.142 ± 0.094 | 2.245 | 1 | 0.134 | 0.574 |
| Germination rate × Style width | −0.040 ± 0.036 | 1.221 | 1 | 0.269 | 0.574 |
| Germination rate × Ovary length | −0.014 ± 0.049 | 0.083 | 1 | 0.773 | 0.918 |
| Germination rate × Ovary width | −0.025 ± 0.046 | 0.285 | 1 | 0.593 | 0.918 |
| **Pollen size** | | | | | |
| Pollen size × Stigma length | −0.051 ± 0.046 | 1.201 | 1 | 0.273 | 0.574 |
| Pollen size × Stigma width | −0.009 ± 0.058 | 0.025 | 1 | 0.875 | 0.918 |
| Pollen size × Stigma area | −0.010 ± 0.058 | 0.029 | 1 | 0.866 | 0.918 |
| Pollen size × Style length | 0.046 ± 0.074 | 0.386 | 1 | 0.534 | 0.918 |
| Pollen size × Style width | −0.009 ± 0.050 | 0.030 | 1 | 0.862 | 0.918 |
| Pollen size × Ovary length | 0.012 ± 0.056 | 0.048 | 1 | 0.827 | 0.918 |
| Pollen size × Ovary width | −0.006 ± 0.062 | 0.011 | 1 | 0.918 | 0.918 |
| **Tube growth rate** | | | | | |
| Tube growth rate × Stigma length | 0.067 ± 0.046 | 2.162 | 1 | 0.141 | 0.574 |
| Tube growth rate × Stigma width | 0.091 ± 0.053 | 2.991 | 1 | 0.084 | 0.574 |
| Tube growth rate × Stigma area | −0.013 ± 0.058 | 0.048 | 1 | 0.826 | 0.918 |
| Tube growth rate × Style length | −0.044 ± 0.066 | 0.425 | 1 | 0.515 | 0.918 |
| Tube growth rate × Style width | 0.132 ± 0.048 | 7.473 | 1 | 0.006 | 0.132 |
| Tube growth rate × Ovary length | 0.086 ± 0.063 | 1.770 | 1 | 0.183 | 0.574 |
| Tube growth rate × Ovary width | 0.158 ± 0.071 | 4.939 | 1 | 0.026 | 0.276 |

Each pollen-pistil trait interaction was tested separately using LRTs comparing models with and without the interaction term. Analyses were based on the complementary experiment, wich included extensive replication of pollen and pistil measurements per parental plant; pollen donor plants contributing less than 5% of fertilized seeds on a given pollen recipient plant were excluded from the analyses. In addition to the pollen-pistil interaction term, single-interaction models included the focal pollen trait, the relative pollen production of competitors as a covariate, and pollen donor plant identity as a random effect. *P*-values were adjusted for multiple testing across all interactions ($n$ = 21) using the FDR method [5]. All pollen and pistil traits were standardized and correspond to predicted values extracted from individual-level random effects of generalized linear mixed-effects models (GLMMs) that corrected for pollen density and other potentially confounding experimental factors included as random effects.
